# Conserved principles of central carbon partitioning in Hippo-Yorkie-driven *Drosophila* gut tumors

**DOI:** 10.64898/2026.05.05.722979

**Authors:** Younshim Park, Mujeeb Qadiri, John M. Asara, Yanhui Hu, Norbert Perrimon

## Abstract

Central carbon metabolism undergoes extensive remodeling in cancers, yet the extent to which the resulting network architectures and operating principles are conserved across species and oncogenic contexts in vivo remains unclear. Here, central carbon metabolism was evaluated in Hippo/Yki-driven *Drosophila* gut tumors, as Hippo-YAP/TAZ signaling links nutritional cues to metabolic state and contributes to epithelial tumorigenesis and therapy resistance. Using integrated steady-state metabolomics, transcriptomics and [U-^13^C_6_]glucose tracing, we defined how Hippo pathway activation reorganizes nutrient utilization and carbon flux in vivo and assessed how the resulting Yki-driven metabolic network aligns with mammalian cancer metabolism. Yki tumors exhibited a Warburg-like state with increased glycolytic throughput and enhanced conversion of glucose-derived carbon to lactate, accompanied by transcriptional upregulation of key glycolytic and lactate-production enzymes. Glucose carbon was also redirected into redox-supporting and anabolic nodes, including activation of the glycerol-3-phosphate shuttle and increased labeling of alanine and serine. Mitochondrial metabolism was reorganized into a non-canonical, segmented TCA network centered on α-ketoglutarate, which accumulated and acted as a drain into glutamate/glutamine and 2-hydroxyglutarate rather than supporting complete oxidative turnover. Despite reduced abundance of pentose phosphate intermediates, non-oxidative PPP carbon rearrangements and ribose labeling were maintained, enabling robust glucose contribution to pyrimidine nucleotide pools, including strongly labeled dTTP. Together, these data establish a comprehensive map of Yki-driven central carbon partitioning in vivo and highlight conserved principles of tumor carbon allocation shared across oncogenic contexts and mammalian cancer metabolism.

## Introduction

Central carbon metabolism integrates nutrient uptake with energy production and biosynthesis, channeling carbon from glucose, glutamine, and other substrates through glycolysis, the tricarboxylic acid (TCA) cycle, and branching anabolic pathways such as the pentose phosphate pathway (PPP)^1,2,3^. These interconnected networks allow cells to tune ATP generation, redox balance, and macromolecular synthesis to changing demands^4^. Although this architectural framework is well established, how carbon flux is dynamically redistributed across these pathways in vivo—and how this redistribution is reprogrammed in disease states—remains incompletely defined.

Oncogenic transformation profoundly remodels central carbon metabolism to sustain growth, support stress resistance, and adapt to fluctuating nutrient environments. Cancer cells commonly exhibit increased glucose uptake and aerobic glycolysis (the Warburg effect), altered TCA cycle engagement, and enhanced PPP activity to generate biosynthetic precursors and maintain NAD(P)H-dependent redox control^5,6^. These metabolic adaptations are actively driven by oncogenic networks, including c-Myc, HIF-1α, mutant KRAS, and the PI3K–AKT–mTOR pathway, which promote glucose and glutamine catabolism and channel carbon into nucleotide, amino-acid, and lipid synthesis^7,8,9,10,11^. However, much of this knowledge derives from mammalian cell culture and mouse models, and it is not clear to what extent the resulting principles and network architectures are conserved across species or oncogenic contexts in vivo.

*Drosophila* has emerged as a powerful system for interrogating tumorigenesis, with neoplastic growth induced by activating oncogenic pathways such as Ras–EGFR, Notch, or Hippo, or by inactivating tumor suppressors including *scribble*, *discs large*, *lethal giant larvae,* and *lats*—genes with clear orthology to mammalian cancer drivers^12,13^. Tumors in these models recapitulate hallmark features of malignancy, including sustained proliferation, resistance to cell death, aberrant nutrient sensing, and invasive behavior^14,15,16,17,18^. Yet, despite this rich genetic toolkit and strong conservation of signaling, relatively few studies have systematically mapped how fly oncogenes and tumor suppressors reorganize central carbon metabolism in vivo or directly compared these programs with canonical metabolic rewiring in mammalian cancers.

The Hippo–YAP/TAZ signaling module is increasingly recognized as a central integrator of mechanical, nutritional, and mitogenic cues with metabolic state, linking tissue growth control to carbon allocation and redox regulation^19,20,21,22^. YAP/TAZ hyperactivation has been implicated in multiple human epithelial cancers and is frequently associated with malignant progression and therapy resistance^23,24,25,26^, yet how Hippo pathway deregulation reshapes metabolic flux at organismal scale remains poorly understood. In *Drosophila*, constitutive activation of Yorkie (Yki)—the homolog of mammalian YAP/TAZ—in the adult intestinal epithelium drives robust tissue overgrowth and recapitulates cardinal features of epithelial tumorigenesis^27,28,29^. How Yki oncogenic signaling orchestrates global reprogramming of central carbon metabolism in these tumors, and whether this rewiring reflects conserved principles of cancer metabolism described in mammals, is unknown.

Here, we leverage a Yki-driven *Drosophila* gut tumor model to define how Hippo pathway activation reorganizes nutrient utilization and carbon flux in vivo. By integrating steady-state metabolomics, transcriptomics, and [U-^13^C₆]glucose tracing, we construct a comprehensive map of how Yki activation coordinately redirects central carbon metabolism during tumor progression. We show that Yki tumors increasingly route glucose-derived carbon away from complete oxidation and toward anabolic biosynthesis. They engage glycolytic branch points, the PPP, and amino-acid and nucleotide synthesis pathways to meet heightened demands for biomass and redox control. Despite this diversion, Yki tumors maintain energy production through aerobic glycolysis supplemented by oxidative phosphorylation. This integrative framework reveals conserved metabolic strategies by which oncogenic signaling rewires central carbon metabolism across species, and highlight *Drosophila* Yki tumors as a tractable platform for dissecting Hippo-dependent metabolic vulnerabilities with direct relevance to YAP/TAZ-activated human cancers.

## Results

### Yki tumors exhibit hallmarks of human cancers at the organismal level

Fly gut tumors were induced by overexpressing an activated Yorkie variant (*yki^3SA^*) in intestinal stem cells using a temperature-sensitive gene-expression driver (Fig. S1A). Immunostaining for phospho-histone H3 (PH3) revealed elevated mitotic activity that was sustained over time (Fig. S1B-D). PH3+ cells increased through day 5 and remained persistently elevated thereafter, consistent with prolonged proliferative activity (Fig. S1B). This persistent proliferative state suggests a requirement for metabolic remodeling to sustain growth, including maintenance of bioenergetic demand, biosynthetic capacity, and redox homeostasis. Consistent with this, differential gene-expression analysis showed significant enrichment of metabolism-associated genes, and metabolite profiling identified broad metabolic remodeling, with prominent changes in macromolecule-related pathways, including amino acids and nucleotide metabolism (Fig. S1I,J).

Day 8 metabolomics, corresponding to an adapted tumor state, further highlighted pathway-level rewiring, with 11 of the 16 most strongly enriched metabolic pathways in Yki tumors also enriched across multiple human cancer datasets (Fig. S2A). Among these, human colon cancer showed the highest concordance. Differential abundance (DA) analysis, which summarizes the net direction of metabolite changes within each pathway, showed concordant up- or downregulation in approximately two-thirds of shared metabolic pathways, including glycolysis and amino acid metabolism (Fig. S2B). This pattern is consistent with partial conservation of tumor metabolic programs.

BODIPY staining revealed reduced lipid stores in tumor-bearing flies, consistent with a cachexia-like systemic alterations in nutrient storage and utilization accompanying tumor growth (Fig. S1E,F)^29^. Moreover, constitutive Yki activation significantly shortened lifespan (Fig. S1G), whereas no survival difference was observed without induction (Fig. S1H), indicating that reduced survival is primarily attributable to tumor burden. Collectively, these data support the Yki gut tumor model for defining central carbon metabolic rewiring in vivo and for comparison with mammalian cancer metabolism; subsequent analyses therefore profiled metabolic remodeling and traced nutrient carbon flow through central carbon pathways.

### Yki-tumors rewire glycolysis to drive a Warburg-like metabolic shift

Under normal conditions, glucose is metabolized through glycolysis to generate pyruvate, which is typically converted into acetyl-CoA (AcCoA) and oxidized in the TCA cycle to support efficient ATP production (Fig. 1A). In contrast, many tumors exhibit aerobic glycolysis (the Warburg effect), preferentially diverting glucose to lactate fermentation despite sufficient oxygen^5^.

**Figure 1.**
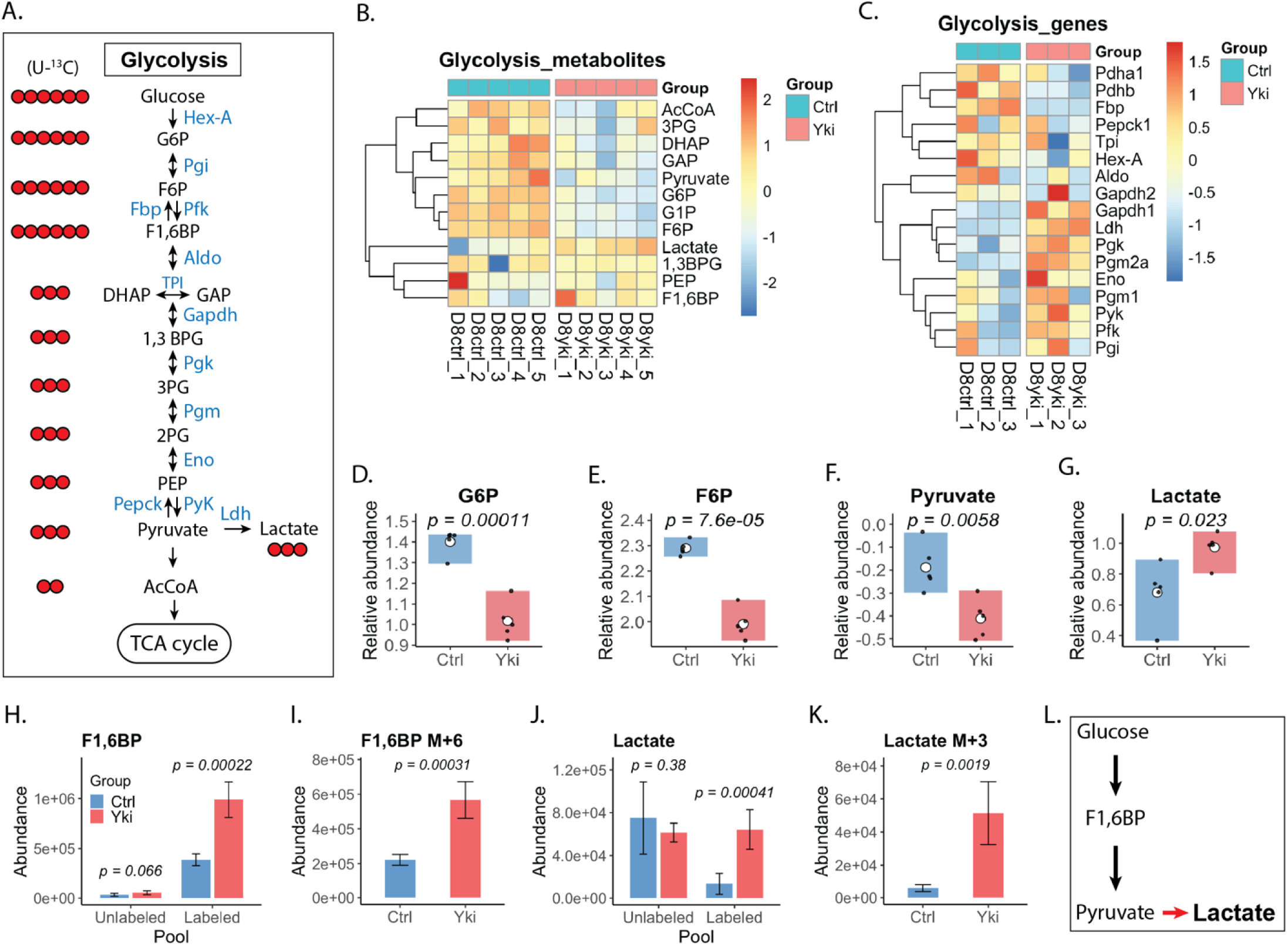
Yki tumors exhibit a Warburg-like metabolic shift. (A) Schematic of glycolysis showing glycolytic enzymes (blue) and intermediates (black). Red circles indicate heavy-isotope ¹³C-labeled glucose isotopologues corresponding to each intermediate. (B) Heatmap of glycolytic metabolite demonstrating broad metabolic changes in Yki tumors. (C) Transcriptomic heatmap indicating altered expression of glycolytic enzymes. (D-G) Stead-state abundance of individual rate-limiting glycolytic metabolites significantly altered in Yki tumors. (H-K) Abundance of ¹³C-labeled glycolytic intermediates shows significant changes, indicating that glycolytic flux is generally increased and redirected toward lactate production. (L) Schematic summary illustrating increased glycolytic flux in Yki tumors and preferential routing of glucose-derived pyruvate toward lactate production. *Statistical comparisons between control and Yki samples were performed using two-tailed t-test. **Heatmap shows scaled values generated using the R package pheatmap. Each row is z-score-normalized across samples to highlight relative differences. Color scale represents row-scaled values, with blue indicating lower and red indicating high relative abundance.

Metabolomic profiling of Yki tumors revealed widespread changes of glycolytic intermediates, with reduced pyruvate and AcCoA alongside elevated lactate, consistent with a Warburg-like metabolic shift (Fig. 1B). Despite depletion of several intermediates, transcripts encoding key glycolytic enzymes—including the rate-limiting *phosphofructokinase* (*Pfk*) and *pyruvate kinase* (*Pyk*)—were increased, suggesting that lower intermediate levels reflect increased glycolytic flux rather than reduced production (Fig. 1C).

Consistent with increased glycolytic commitment, the upstream intermediates glucose-6-phosphate (G6P) and fructose-6-phosphate (F6P) were markedly depleted in Yki tumors, suggesting accelerated Pfk-driven consumption (Fig. 1D,E). [U-¹³C₆]glucose tracing further supported enhanced glycolytic engagement, with increased total abundance and labeled fraction of fructose-1,6-bisphosphate (F1,6BP) in Yki tumors, indicating elevated incorporation of glucose-derived carbon at this key glycolytic node (Fig. 1H,I).

Downstream metabolites showed a concordant labeling pattern. The labeled fraction of dihydroxyacetone phosphate (DHAP), glyceraldehyde-3-phosphate (GAP), and phosphoenolpyruvate (PEP) were significantly increased in Yki tumors, consistent with enhanced propagation of glucose-derived carbon through lower glycolysis (Fig. S3D-F). In contrast, pyruvate abundance was reduced despite increased *Pyk* expression and evidence of elevated upstream flux (Fig. 1F). Given the rapid turnover of pyruvate, this pattern is consistent with increased downstream utilization rather than reduced production.

Lactate measurements further supported altered metabolic fate in Yki tumors. Lactate abundance was substantially increased relative to controls (Fig. 1G). Consistent with this increase, [U-¹³C₆]glucose tracing showed enrichment of the lactate M+3 isotopologue with a corresponding reduction in the unlabeled (M+0) fraction (Fig. 1J,K; Fig. S3H), indicating increased incorporation of glucose-derived carbon into lactate and enhanced pyruvate-to-lactate conversion. *Lactate dehydrogenase (Ldh)* expression was up regulated in Yki tumors, consistent with transcriptional reinforcement of elevated lactate production ((Fig. S3I). Together, these data indicate increased glycolytic flux in Yki tumors that is preferentially directed toward lactate production, consistent with a Warburg-like glycolytic program (Fig. 1L).

### Yki tumors rewire the G3P shuttle for compartmentalized redox control and potential lipid biosynthesis

Enhanced glycolysis can support rapid ATP generation and provide intermediates for nucleotide, lipid and amino acid biosynthesis^7,30,31^. The glycerol-3-phosphate (G3P) shuttle couples cytosolic glycolysis to mitochondrial metabolism and also supplies G3P for lipid synthesis; this axis is frequently remodeled in cancers in the context of Warburg-like metabolism^32,33^ (Fig. 2A). In Yki tumors, steady-state G3P abundance was unchanged (Fig. 2B), whereas [U-¹³C₆]glucose tracing showed significantly increased glucose-derived labeling of G3P (Fig. 2D), with enrichment of the G3P M+2 and M+3 isotopologues (Fig. 2E; S4A). Fractional labeling further showed increased G3P M+3 with a reciprocal decrease in the unlabeled (M+0) fraction demonstrating increased diversion of glucose-derived carbon into this node (Fig. 2F). Consistent with activation of the shuttle, *Gpo1* (mitochondrial glycerol-3-phosphate dehydrogenase) transcripts were markedly increased, suggesting that Yki tumors recruit this pathway to meet their metabolic demands (Fig. 2C; S4J).

**Figure 2.**
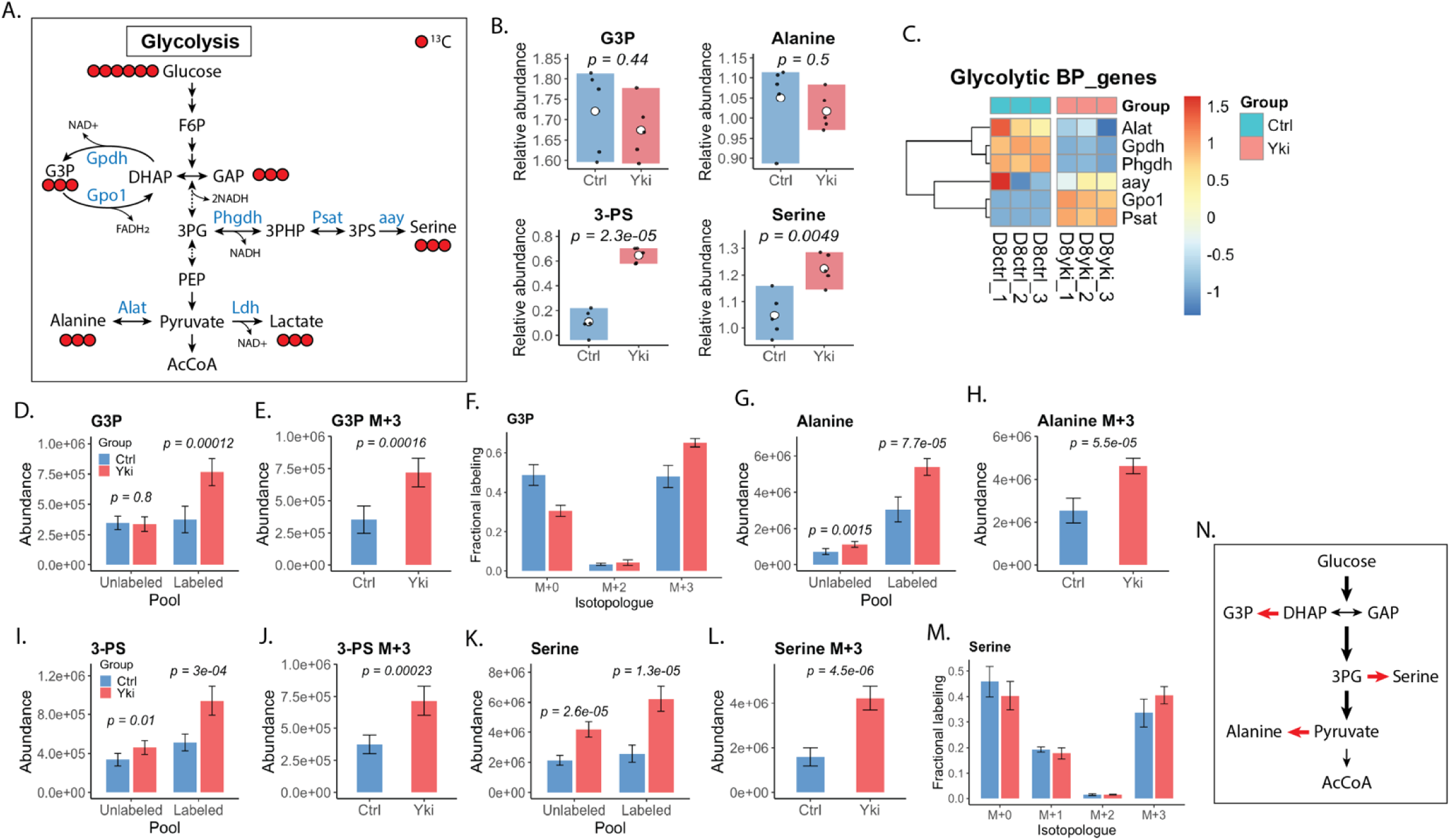
Yki tumors rewire glycolytic branch points to drive the glycerol-3-phosphate shuttle and alanine/serine biosynthesis. (A) Schematic of glycolysis highlighting the glycerol 3-phosphate (G3P) shuttle and branch points into alanine and serine biosynthesis. Red circles indicate ¹³C atoms derived from [U-¹³C₆]glucose; enzymes quantified in this study are indicated (Gpdh, Gpo1, Alat, Phgdh, Psat, aay; Ldh shown for reference). (B) Relative abundances of G3P, alanine, 3-phosphoserine (3-PS), and serine in control (Ctrl) and Yki tumor (Yki) samples. (C) Heatmap of glycolytic branch-point (BP) genes involved in G3P, alanine, and serine metabolism showing scaled expression values from RNA-seq for individual Ctrl and Yki samples. (D–F) [U-¹³C₆]glucose tracing of G3P. (D) Abundances of unlabeled (M+0) and total ¹³C-labeled (ΣM+n) G3P pools. (E) Abundance of the fully labeled G3P isotopologue (M+3). (F) Fractional labeling of G3P isotopologues (M+0, M+2, M+3). (G, H) [U-¹³C₆]glucose tracing of alanine. (G) Abundances of unlabeled and total ¹³C -labeled alanine pools. (H) Abundance of alanine M+3. (I, J) [U-¹³C₆]glucose tracing of 3-PS. (I) Abundances of unlabeled and total ¹³C-labeled 3-PS pools. (J) Abundance of 3-PS M+3. (K, L, M) [U-¹³C₆]glucose tracing of serine. (K) Abundances of unlabeled and total ¹³C-labeled serine pools. (L) Abundance of serine M+3. (M) Fractional labeling of serine isotopologues (M+0, M+1, M+2, M+3).(N) Schematic summary of glucose carbon partitioning in Yki tumors. Glucose-derived carbon is routed through the G3P node in exchange with DHAP and GAP, and is redirected from glycolytic intermediates toward anabolic outputs. Red arrows indicate pathways showing increased glucose contribution in Yki tumors, including diversion from DHAP to G3P, 3PG to serine and from pyruvate to alanine.

Despite increased G3P labeling, *Gpdh* (cytosolic glycerol-3-phosphate dehydrogenase) expression and total DHAP abundance were reduced (Fig. S3B; Fig. 2C; Fig. S4I), arguing against bulk substrate accumulation as the primary driver of flux. Isotopologue analysis resolved this discrepancy: although the DHAP pool was dominated by M+2 (Fig. S3G), Yki tumors showed selective enrichment of glucose-derived M+3 DHAP and near-exclusive propagation of this M+3 label into G3P (Fig. S3E; 2E,F). Together these labeling relationships indicate preferential use of a newly synthesized glycolytic DHAP M+3 pool for G3P production rather than the PPP-associated M+2 pool, consistent with substrate channeling into the G3P shuttle and with mitochondrial Gpo1 serving as a downstream sink to sustain flux.

Routing DHAP into G3P also provides a mechanism for redox control, as DHAP-to-G3P conversion consumes NADH and regenerates NAD+, complementing lactate dehydrogenase-mediate NAD+ regeneration (Fig. 2A)^32^. Consistent with increased NADH-consuming capacity, Yki tumors showed increased NAD+ and decreased NADH, with a trend toward an increased NAD+/NADH ratio (Fig. S4K,L). The combination of strong activation of NADH-consuming pathways with modest changes in the bulk ratio is consistent with redox compartmentalization, in which a high cytosolic NAD+ pool supporting glycolysis is partially masked by more buffered mitochondrial pools in whole-tissue measurements.

Beyond redox homeostasis, G3P is an entry point to glycerolipid and glycerophospholipid synthesis, and a large portion of genes linked to phospholipid metabolism were coordinately upregulated in Yki tumors (Fig. S4M). Together with the increased diversion of glucose-derived carbon into G3P, these data support a model in which Yki tumors expand G3P-centered metabolism to meet anabolic demands, with potential conservation relative to human cancers. Direct measurements of lipid synthetic flux will be required to establish the extent to which G3P is routed into phospholipid production in this model.

### Enhanced glycolysis channels glucose carbons into alanine and serine biosynthesis in Yki tumors

In mammalian cancers, elevated glycolysis is also frequently coupled to anabolic amino acid production^34,35^. Glycolytic intermediates can be diverted to de novo serine (and downstream glycine) synthesis via 3-phosphoglycerate (3-PG), and to alanine synthesis via transamination of pyruvate; both programs are recurrently upregulated across diverse human tumors^35,36^ (Fig. 2A). In Yki tumors, metabolomics and [U-¹³C₆]glucose tracing showed significant increases in 3-phosphoserine (3-PS) and serine whereas total alanine abundance was unchanged (Fig. 2B). Despite stable alanine levels, ^13^C-labeled alanine, 3-PS, and serine were increased (Fig. 2G,I,K), with strong enrichment of the M+3 isotopologue for each metabolite (Fig. 2H,J,L), indicating diversion of glucose-derived carbon from pyruvate and 3-PG into amino acid biosynthesis.

Alanine production appeared to increase at the level of flux rather than steady-state concentration. Both unlabeled and ¹³C-labeled alanine pools were higher in Yki tumors, and alanine M+3 was markedly increased, consistent with elevated transamination of glycolytic pyruvate (Fig. 2G,H; S4B-D). In contrast, the alanine transaminase transcript was reduced (Fig. 2C), suggesting that increased alanine labeling is driven predominantly by substrate availability and/or post-transcriptional regulation rather than transcriptional induction. This combination of increased glucose-derived labeling with stable bulk abundance suggests tight regulatory constraints on pyruvate partitioning, although the downstream utilization of alanine carbon remains to be defined. This configuration also differs from several mammalian tumors in which de novo alanine synthesis is reduced, and alanine supply is instead maintained via stromal export^36,37^, indicating context-dependent modes of alanine metabolism.

By contrast, the serine branch exhibited a canonical cancer-like remodeling. Yki tumors accumulated 3-PS and serine (Fig. 2B) and showed significant increases in both unlabeled and ¹³C-labeled pools, with a pronounced rise in serine M+3 (Fig. 2I–L; S4E-H). Fractional labeling showed enrichment of serine M+3 with a concomitant reduction in the unlabeled (M+0) fraction (Fig. 2M), indicating incorporation of glucose-derived carbon into de novo serine synthesis. The unlabeled serine pool was also increased in Yki tumors, consistent with overall expansion of the serine pool (Fig. 2K). However, the increased M+3 fraction and the decreased M+0 fraction indicate that a larger share of this expanded pool is supplied by de novo synthesis from glucose (Fig. 2M). The remaining unlabeled component likely reflects continued uptake of exogenous serine and/or production from non-glucose precursors, including reversible serine-glycine interconversion via serine hydroxymethyltransferase (Shmt) and one-carbon metabolism (Fig. S4N).

Given that serine supplies glycine and one-carbon units to support nucleotide and methionine metabolism (Fig. S4N-T), increased serine provides a potential route for glucose carbon to contribute to the elevated M+1 methionine observed in Yki tumors, consistent with folate-dependent one-carbon transfer (Fig. S4U,V). In parallel, increased M+2 and M+3 methionine isotopologues suggest uptake of ¹³C-labeled methionine from the microenvironment, potentially produced by microbiota from dietary glucose (Fig. S4W,X). Dissecting the relative contributions of one-carbon-dependent methionine regeneration versus exogenous will require dedicated flux experiments, such as parallel tracing with ¹³C-methionine and microbiota-modulating conditions. Together, these data indicate that activation of the serine synthesis represents a conserved cancer metabolic phenotype shared between Yki tumors and mammalian cancers, whereas the distinct regulation of alanine labeling highlights divergence in glycolytic carbon partitioning across tumor contexts and species (Fig. 2N).

### Yki tumors display a non-canonical, “segmented” TCA cycle centered on α-ketoglutarate

Given the increased glycolytic flux and diversion of glucose carbon into anabolic pathways, mitochondrial carbon handling was next examined. Whereas differentiated tissues typically prioritize oxidative TCA cycling for efficient ATP production (Fig. 3A), proliferating cells often divert TCA intermediates to support biosynthesis^34,38,39^. In Yki tumors, metabolomic profiling revealed marked reorganization of the TCA network (Fig. 3B). Abundances of early intermediates (AcCoA, citrate, cis-aconitate (cisAcon) and isocitrate) and the downstream intermediate succinate were significantly depleted (Fig. 3D–G,I), whereas α-ketoglutarate (α-KG) was consistently elevated (Fig. 3H). Transcriptomic analysis showed broad downregulation of TCA cycle genes, with relative preservation of *Idh* and *Scs* transcripts encoding isocitrate dehydrogenase and succinyl-CoA synthetase, respectively (Fig. 3). Together, these data indicate a localized metabolic imbalance centered on the α-KG node.

**Figure 3.**
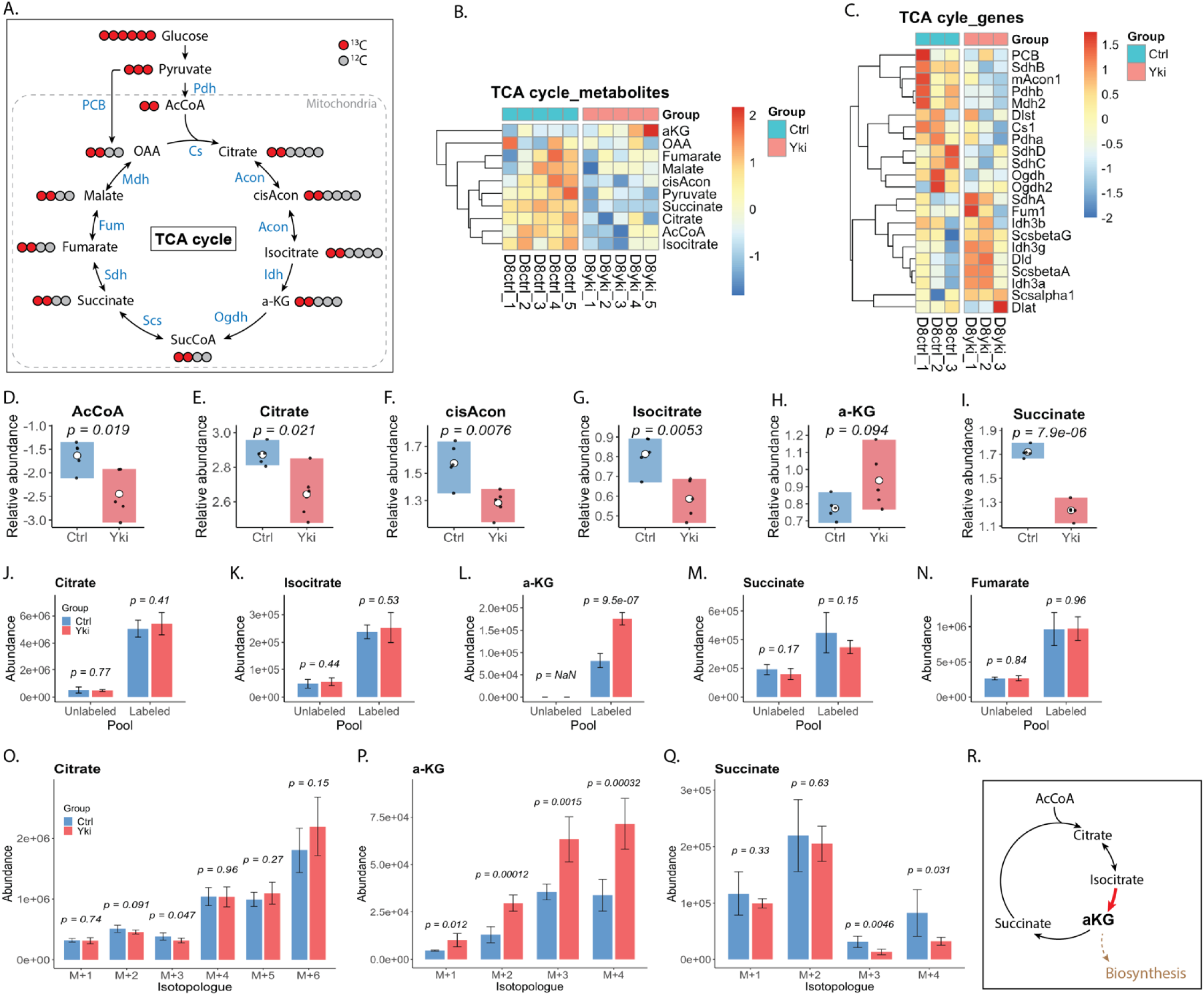
α-ketoglutarate as a bottleneck in the Yki tumor TCA network. (A) Schematic of the TCA cycle and associated anaplerotic entry points illustrating the complete oxidation of glucose-derived carbon and the generation of TCA intermediates within mitochondria. Red circles denote the primary carbon flow from glycolytic-derived, labeled intermediates. (B) Heatmap of TCA-cycle metabolite abundances showing broad remodeling of central carbon metabolism in Yki tumors, with a prominent shift at the α-KG node. (C) Heatmap of TCA-cycle gene expression indicating coordinated downregulation of multiple TCA enzyme complexes, with relative preservation of genes involved in α-KG production in Yki tumors. (D–I) Relative abundance of individual TCA intermediates. Early intermediates (AcCoA, citrate, cis-aconitate, isocitrate) and the downstream intermediate succinate are depleted, whereas α-KG is increased in Yki tumors. (J–N) Abundance of unlabeled and ¹³C-labeled pools for representative intermediates (citrate, isocitrate, α-KG, succinate and fumarate), highlighting preserved glucose contribution to most intermediates but a disproportionately increased ¹³C-labeled α-KG pool. (N) Citrate isotopologue abundances (M+1–M+6), illustrating enrichment of higher-order labeled species consistent with extensive incorporation and cycling of glucose-derived carbon. (O) α-KG isotopologue abundances (M+1–M+4) showing increased labeled species across the distribution in Yki tumors. (P) Succinate isotopologue abundances (M+1–M+4) showing attenuated propagation of higher-order label relative to α-KG, consistent with an isotopic discontinuity downstream of α-KG. (Q) Schematic summary depicting a segmented TCA organization in Yki tumors, with α-KG positioned as a bottleneck/branch point that diverts carbon toward biosynthetic fates. *Bars show mean ± s.d.; points indicate biological replicates. For isotopologue panels (Fig. 3O–Q), statistical comparisons between Ctrl and Yki were performed using Welch’s two-sided t-test (unequal variances), applied independently to each isotopologue (e.g., M+0, M+1, M+2, M+3, etc.); exact P values are shown on the plots.

To distinguish altered synthesis from altered turnover, [U-¹³C₆]glucose tracing was used to assess glucose contribution to TCA pools. Despite reduced abundance, citrate and isocitrate showed comparable ¹³C fractional enrichment in control and Yki tumors, with glucose-derived carbon remaining the dominant contributor (Fig. 3J,K,O; Fig. S5A). Downstream intermediates (succinate, fumarate, malate, and oxaloacetate (OAA)) similarly showed largely preserved ¹³C-labeling (Fig. 3M,N,Q; Fig. S5>D–I). In contrast, α-KG exhibited a distinct signature, with markedly increased ¹³C incorporation and a minimal unlabeled fraction, indicating that the expanded α-KG pool is sustained predominantly by glucose-derived carbon (Fig. 3L,P). These patterns suggest increased turnover or efflux for most TCA intermediates, alongside selectively enhanced input into and/or restricted utilization at the a-KG node.

Isotopologue distributions further resolved carbon flow through the pathway. Citrate M+2 primarily reports canonical oxidative entry of [U-¹³C₆]glucose via pyruvate and acetyl-CoA, whereas higher-order isotopologues (M+3–M+6) indicate multiple turns of the cycle and anaplerotic inputs such as pyruvate-to-oxaloacetate conversion. Citrate showed substantial higher-order isotopologues, consistent with extensive incorporation and cycling of glucose-derived carbon and anaplerotic input (Fig. 3O; S5A). Among TCA intermediates, α-KG showed the strongest shift: in Yki tumors, all labeled α-KG isotopologues (M+1–M+4) were significantly increased, and α -KG fractional enrichment exceeded that of upstream citrate and downstream succinate, malate, and OAA (Fig. 3O–Q; Fig. S5F-I). This localized accumulation and hyper-labeling, without comparable enrichment in flanking metabolites, is consistent with a metabolic bottleneck at the α-KG.

An isotopic discontinuity between citrate/ α-KG and succinate provided additional evidence for “segmentation” of the cycle. Although citrate exhibited low unlabeled (M+0) fraction and abundant higher-order labeled species (M+3–M+6) (Fig. 3O; Fig. S5A), succinate retained a dominant unlabeled fraction and showed significantly reduced incorporation of the M+3/M+4 species that were abundant in α-KG (Fig. 3P,Q). This dissociation suggests that the glucose-derived carbon entering the early TCA cycle does not efficiently propagate through α-KG dehydrogenase into downstream intermediates, consistent with diversion of carbon away from complete oxidative turnover. Collectively, these data support a non-canonical, “segmented” TCA organization in Yki tumors, with α-KG functioning as a central hub for carbon rerouting (Fig. 3R).

### α-ketoglutarate functions as a carbon drain point in the Yki tumor TCA network

In mammalian cancers, α-KG links TCA-cycle activity to amino-acid biosynthesis and to production of 2-hydroxyglutarate (2-HG), a metabolite with implicated in epigenetic regulation^40,41,42,43^. Given the segmented TCA organization and selective accumulation of α-KG in Yki tumors (Fig. 3), glucose-derived carbon was hypothesized to be preferentially diverted into α-KG-dependent biosynthetic branches (Fig. S5K).

Consistent with this model, [U-¹³C₆]glucose tracing showed that products downstream of α-KG-connected closely tracked α-KG labeling. Glutamate and glutamine, which direct derive directly from the α-KG carbon skeleton, maintained steady-state pool sizes (Fig. S5L) yet exhibited significantly increased incorporation of glucose-derived carbon across all detectable isotopologues in Yki tumors (Fig. S5N-P, R-T). This shift was accompanied by increased transcript abundance for enzymes mediating α-KG exit routes, including aminotransferases and glutamine synthetase (*Gs*) (Fig. S5M). In parallel, 2-HG showed increased steady-state abundance (Fig. S4L) and elevated ¹³C isotopologue abundances in Yki tumors (Fig. S5Q,U). Together, these data indicate that the α-KG node functions as a major carbon drain, redirecting glucose-derived carbon from the truncated TCA cycle into glutamate/glutamine biosynthesis and 2-HG production rather than complete oxidative turnover.

Despite overall concordance between α-KG and its downstream products, analysis of fully labeled species revealed a marked discordance. The α-KG M+5 isotopologue was below the limit of detection in both control and Yki tissues (Fig. S5B), whereas glutamate and glutamine contained substantial M+5 fractions that were further increased in Yki tumors (Fig. S5R,S). Because M+5 glutamate/glutamine requires fully labeled α-KG, these data imply transient formation of α-KG M+5 that does not accumulate in the bulk α-KG pool. This pattern is consistent with a small, rapidly turning-over sub-pool that is tightly coupled to amino-acid synthesis. Notably, M+5 labeling was prominent in glutamate and glutamine but comparatively low in 2-HG despite robust overall 2-HG labeling (Fig. S5U), suggesting preferential routing of this highly labeled α-KG sub-pool toward nitrogen metabolism.

Collectively, these observations support a model in which Yki tumors generate a privileged “biosynthetic” α-KG micro-pool—potentially defined by subcellular compartmentation and/or enzyme-proximal coupling—that is rapidly replenished by glucose-derived carbon and consumed by aminotransferases, glutamate dehydrogenase, and glutamine synthetase before equilibrating with the larger, heterogeneous α-KG pool detected by bulk metabolomics. During the tracing period, rapid turnover of this micro-pool can yield substantial M+5 glutamate and glutamine, whereas M+2–M+4 species likely reflect contributions from larger, more dilute α-KG pools shared with residual oxidative TCA reactions and other pathways. Together, these data indicate that Yki activation rewires carbon partitioning at the α-KG node to favor anabolic outputs from glucose over complete oxidation (Fig. S5J,V).

### Non-oxidative PPP enhances glucose carbon channeling in Yki tumors

In addition to diversion of glucose carbon through the α-KG node, glucose-derived intermediates can be repartitioned through the pentose phosphate pathway (PPP) to support redox balance and anabolism. The oxidative PPP (ox-PPP) produces NADPH and ribose-5-phosphate (R5P), whereas the non-oxidative PPP (non-oxPPP) interconverts glycolytic intermediates to generate precursors for nucleotide and amino acid biosynthesis (Fig. 4A)^44,45,46,47^.

**Figure 4.**
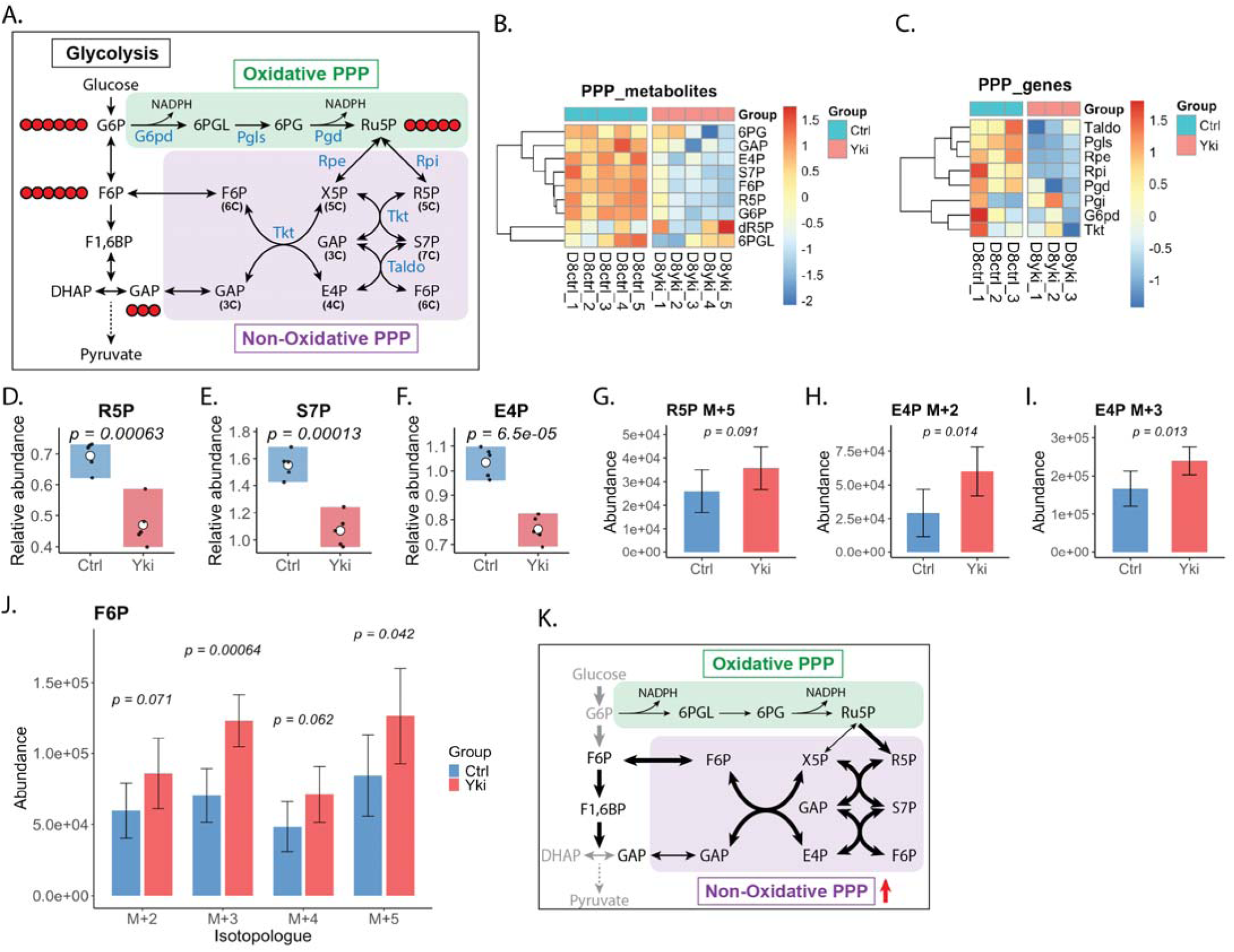
Yki tumors channel glucose carbon via the non-oxidative PPP. (A) Schematic of the pentose phosphate pathway (PPP) illustrating carbon flow from glycolysis into the oxidative (ox-PPP) and non-oxidative (non-oxPPP) branches. (B) Heatmap of PPP metabolites showing broadly decreased levels of pathway intermediates. (C) Transcriptomic heatmap demonstrating widespread downregulation of PPP enzyme expression at the mRNA level. (D–F) Steady-state abundances of PPP intermediates are significantly decreased in Yki tumors. (G–I) Abundance of ¹³C-labeled metabolites indicating increased carbon flow into the non-oxPPP. (J) Abundance of ¹³C-labeled F6P isotopologues supports tight channeling between glycolysis and the non-oxPPP. (K) Schematic summary of the PPP in Yki tumors, highlighting increased glucose-derived carbon channeling between glycolytic intermediates and the non-oxPPP.

In Yki tumors, metabolomics and transcriptomics showed broad depletion of PPP intermediates accompanied by downregulation of PPP-associated genes, consistent with overall pathway suppression at both metabolite and transcriptional levels (Fig. 4B,C). However, [U-¹³C₆]glucose tracing uncovered a distinct pattern: the R5P M+5 isotopologue, which reports direct entry of glucose carbon through ox-PPP, was moderately increased despite a reduced total R5P pool (Fig. 4D,G). This indicates that ox-PPP activity is constrained but remains active. Consistent with allosteric regulation at the level of cofactor availability, NADP+ abundance was significantly decreased in Yki tumors (Fig. S6F), which could limit G6PD activity and ox-PPP throughput, yet ¹³C-labeled R5P production persisted under these conditions (Fig. S6A-C).

Notably, although steady-state levels of non-oxPPP intermediates such as sedoheptulose-7-phosphate (S7P) and erythrose-4-phosphate (E4P) were reduced, labeling analysis showed increased ¹³C-labeled E4P (M+2/M+3) and increased labeling of fructose-6-phosphate (F6P; M+2–M+5) in Yki tumors (Fig. 4H–J, S6E). These patterns support increased non-oxPPP carbon shuffling that is not explained by transcript upregulation, consistent with flux control dominated by metabolic allostery. In parallel, NADPH abundance and NADP+/NADPH ratio were maintained despite low NADP+, suggesting that NADPH homeostasis is supported by sources beyond the ox-PPP (Fig. S6F,G).

Collectively, these data indicate that ox-PPP activity in Yki tumors is constrained, likely by limited NADP+ availability, whereas the non-oxPPP adopts a dominant role in redistributing glucose carbon into biosynthetic precursors. This organization is consistent with reprogramming of central carbon metabolism that sustains anabolic demands despite restricted ox-PPP capacity.

### Yki tumors exhibit enhanced glucose contribution to pyrimidine nucleotide synthesis

Given increased non-oxidative PPP carbon shuttling in Yki tumors and the close coupling of the PPP to nucleotide metabolism, nucleotide biosynthesis was next examined. Most tissues rely on intracellular de novo synthesis and salvage to meet nucleotide demand, as dietary nucleic acids minimally contribute to cellular nucleotide pools^48,49,50^. Rapidly proliferating tumors commonly increase nucleotide production from central carbon metabolism: elevated glucose uptake and glycolytic flux support nucleotide synthesis by providing ribose-5-phosphate (R5P) through the pentose phosphate pathway (PPP) and coordinating inputs from glutamine, aspartate, and serine/one-carbon metabolism for purine and pyrimidine ring assembly (Fig. 5A; Fig. S7G)^51,52,53^. Notably, the non-oxPPP can represent a major route for ribose production in tumors, underscoring the importance of glycolysis-to-PPP carbon rearrangements in supporting nucleotide biosynthesis^54^.

**Figure 5.**
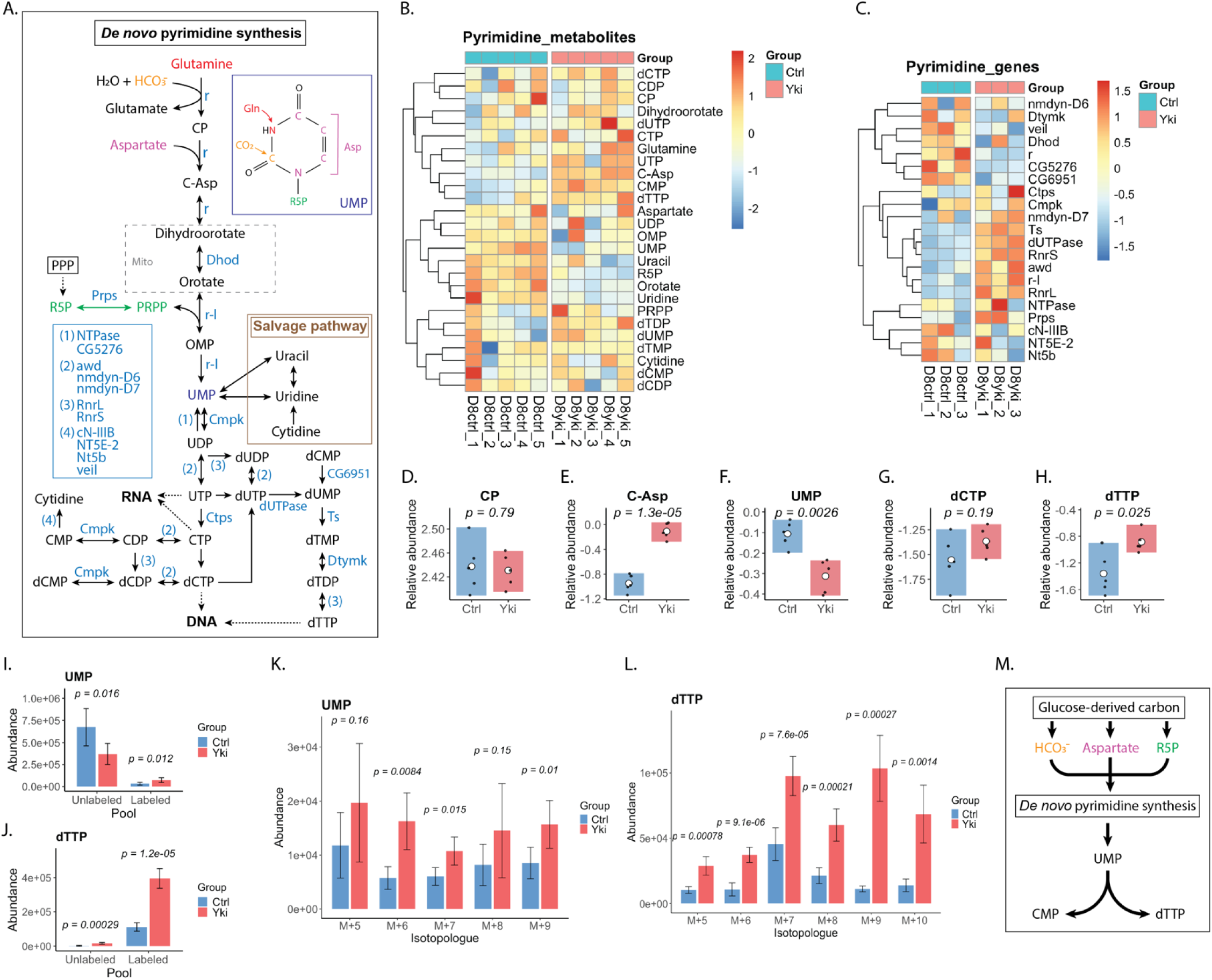
De novo pyrimidine synthesis and salvage are remodeled in Yki tumors, with altered dTTP pools. (A) Schematic of de novo pyrimidine synthesis and the uracil/uridine salvage branch. Enzymes are shown in blue and metabolites in black. Colored substrates indicate major inputs into pyrimidine production (e.g., glutamine-, HCO₃⁻-, aspartate-, and ribose/PRPP-derived contributions) that ultimately support RNA- and DNA-directed nucleotide pools. (B) Heatmap of pyrimidine-pathway metabolite abundances, showing pathway-wide shifts in Yki tumors. (C) Heatmap of pyrimidine-pathway gene expression, indicating coordinated transcriptional remodeling of biosynthetic, interconversion, and salvage-associated enzymes.(D–H) Relative abundance of representative intermediates and products (carbamoyl phosphate, carbamoyl aspartate, UMP, dCTP, and dTTP), highlighting selective changes across the pathway. (I–J) Abundance of unlabeled and ¹³C-labeled pools for UMP and dTTP following [U-¹³C₆]glucose tracing, reporting glucose contribution to ribose-containing nucleotide pools. (K) UMP isotopologue abundances (M+5–M+9), quantifying glucose-derived ¹³C-labeling of the ribose and total carbon backbone. (L) dTTP isotopologue abundances (M+5–M+10), quantifying incorporation of glucose-derived carbon into deoxythymidine triphosphate. (M) Schematic summary highlighting glucose-supported inputs (HCO₃⁻, aspartate, and ribose pathway intermediates) that converge on de novo pyrimidine synthesis to supply UMP and downstream CMP/dTTP pools. *Bars show mean ± s.d.; points indicate biological replicates. Statistical comparisons between Ctrl and Yki were performed using Welch’s two-sided t-test (unequal variances). For isotopologue panels, Welch’s two-sided t-test was applied independently to each isotopologue (e.g., M+0, M+1, M+2, etc.); exact P values are shown on the plots. Metabolites that appear in multiple pathways (e.g., glutamine) are shown in each relevant heatmap.

**Figure 6.**
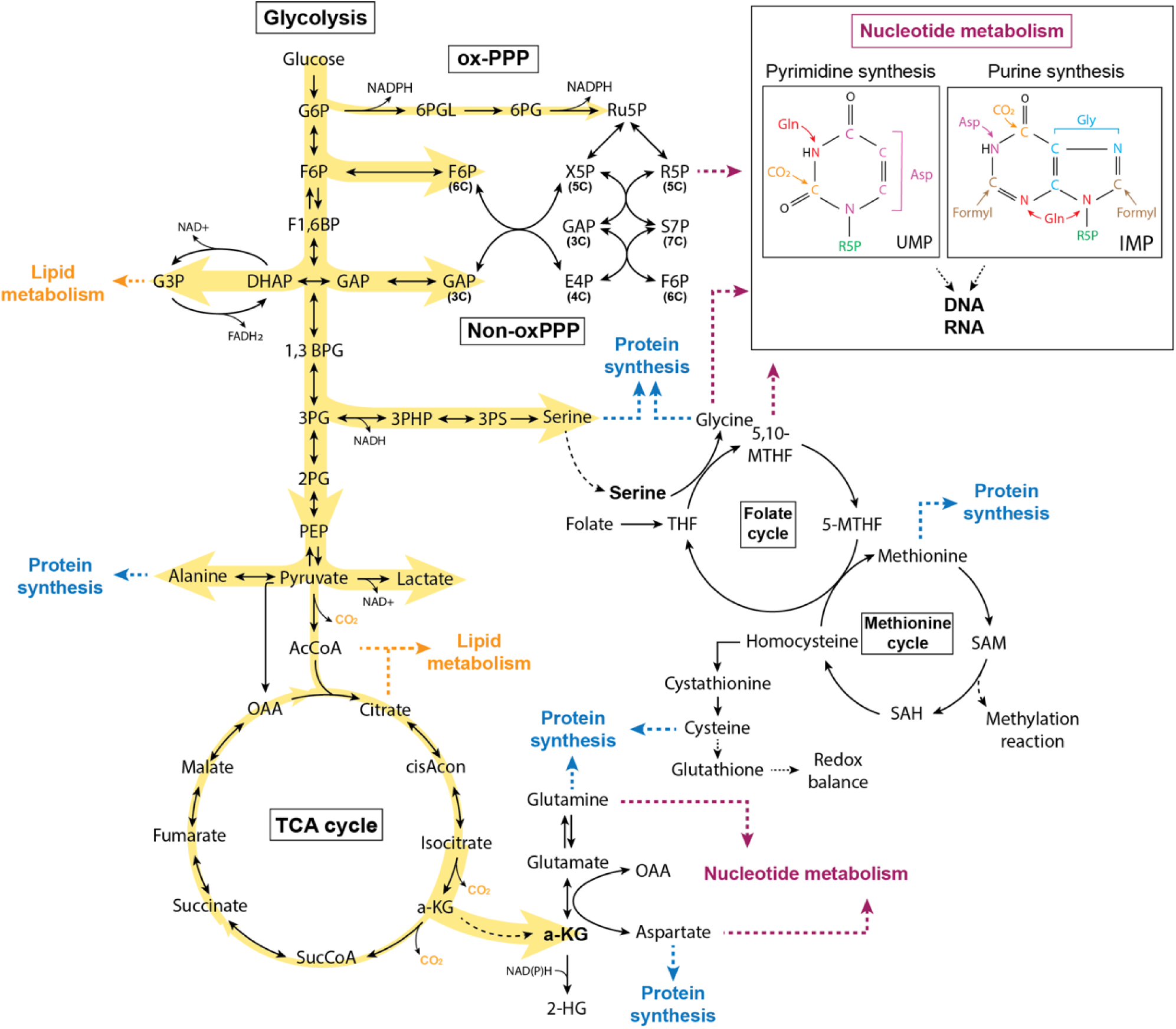
Schematic of glucose-derived carbon fate in Yki tumors. Yki tumors rewire central carbon metabolism by enhancing glycolytic flux and diverting glucose-derived carbon away from complete mitochondrial oxidation toward redox balance and biosynthetic production.

Consistent with this framework, day 8 (D8) Yki tumors showed enhanced glycolysis, increased non-oxPPP carbon shuffling with moderately increased labeling of R5P, and diversion of glucose-derived carbon into serine/glutamate/glutamine/aspartate, collectively supporting increased routing of glucose carbon into nucleotide-supporting pathways^52,55^. Nucleotide metabolism was broadly remodeled at both metabolite and transcript levels, with prominent changes in de novo pyrimidine synthesis (Fig. 5B,C; Fig. S7H). Among early pyrimidine intermediates, carbamoyl-aspartate (C-Asp) accumulated whereas orotate was reduced; concordantly, *Dhod* expression (encoding dihydroorotate dehydrogenase, DHODH) was decreased, consistent with reduced capacity at the dihydroorotate-to-orotate step (Fig. 5B-E). Despite these upstream alterations, downstream deoxynucleotide pools were elevated, including a significant increases in dTTP (Fig. 5H). Similarly, dATP abundance was also significantly increased in purine synthesis (Fig. S7H,K).

[U-¹³C₆]glucose tracing further resolved glucose contribution to nucleotide pools. C-Asp labeling was unchanged, whereas PRPP labeling increased modestly (Fig. S7A,B), consistent with increased ribose pathway input. Glucose-derived carbon incorporation was significantly increased in UMP, a node supplied by both de novo synthesis and uridine salvage that feeds downstream pyrimidine nucleotide production (Fig. 5K), reflected by decreased unlabeled UMP and increased ^13^C-labeled isotopologues (Fig. 5I). Downstream pyrimidine nucleotides (including UTP and CMP) similarly showed increased glucose-derived labeling (Fig. S7C-F). Strikingly, dTTP exhibited a pronounced shift toward the labeled pool, with ^13^C-labeled isotopologues predominating over unlabeled (Fig. 5J,L). In contrast, dATP accumulation was driven primarily by the unlabeled pool, with little change in labeled dATP (Fig. S7K), whereas dGTP showed increased labeling despite an unlabeled-dominant pool (Fig. S7L,N).

Together, these data indicate expanded dNTP pools with distinct substrate contributions for pyrimidines versus purines in Yki tumors. The coordinated increase in labeled UMP/UTP/CMP and the predominance of labeled dTTP support preferential routing of glucose-derived carbon into pyrimidine nucleotide synthesis and onward into the dTTP pool (Fig. 5M). Whether comparatively weaker labeling of purine dNTPs reflects altered purine turnover rather than differential biosynthetic routing remains to be determined. Finally, C-Asp accumulation together with reduced orotate and lower *Dhod* expression suggests a potential constraint at, or downstream of, the dihydroorotate–orotate node. Nevertheless, downstream pyrimidine nucleotides remained elevated and strongly labeled, consistent with compensation via increased downstream demand, salvage contributions and/or altered nucleotide turnover. Because metabolite abundances and isotopologue distributions alone cannot quantify pathway flux or deconvolve de novo versus salvage contributions, resolving the dominant route will require complementary tracer strategies and/or formal metabolic flux analysis.

## Discussion

This study delineates a Hippo/Yki-driven metabolic program in adult *Drosophila* gut tumors. Tumorigenesis rewires central carbon metabolism by enhancing glycolytic flux and diverting glucose-derived carbon away from complete mitochondrial oxidation toward NAD⁺-dependent redox maintenance and biomass production. Integration of metabolomics, transcriptomics and [U-¹³C₆]glucose tracing indicates that Yki tumors adopt an in vivo Warburg-like metabolic state while coordinately reconfiguring multiple nodes of central carbon metabolism. Collectively, these changes support a growth-associated organization in which glucose is funneled through glycolysis, redox-regenerating pathways sustain that flux, mitochondrial oxidation is curtailed and carbon is redistributed into biosynthetic outputs that directly support proliferation.

Mechanistically, Yki tumors display increased ¹³C labeling of committed and downstream glycolytic intermediates together with elevated transcripts of the rate-limiting enzymes Pfk and Pyk, consistent with increased glycolytic throughput. Strong enrichment of ¹³C-labeled lactate and glycerol-3-phosphate, accompanied by upregulation of *Ldh* and the mitochondrial G3P dehydrogenase *Gpo1*, indicates preferential routing of glucose-derived carbon into these downstream fates in parallel with augmented NAD⁺ regeneration. Glucose-derived carbon is also directed into anabolic pathways: glycolytic intermediates feed alanine and serine biosynthesis via pyruvate and 3-PG, respectively, and within the TCA network α-KG emerges as a major drain node into glutamate, glutamine and 2-HG. In addition, the PPP exhibits increased non-oxidative carbon rearrangements, with robust glucose contribution to pyrimidine nucleotides (UMP, UTP and CMP) and increased labeling of dTTP. Together, these features indicate that Yki tumors use glucose as a carbon source for redox support and for synthesis of amino acids and nucleotides, rather than as a substrate for maximal oxidative ATP production.

Multi-omic profiling of Yki-driven gut tumors recapitulates core features of central carbon remodeling described across mammalian cancers. Increased glycolytic throughput with a Warburg-like shift toward lactate production is observed across malignancies, including triple-negative breast cancer, pancreatic cancer, lung cancer and colorectal cancer^56,57,58,59,60,61,62^, and Yki tumors show a corresponding combination of elevated glycolytic labeling, increased lactate abundance and *Ldh* expression. Downstream of glycolysis, Yki tumors also engage the serine–one-carbon axis in a cancer-like manner: serine abundance and glucose-derived serine labeling are increased, and serine pathway enzymes are upregulated, consistent with mammalian programs in which glucose-derived serine supports proliferation and stress resistance^63,64,65^. Notably, whereas several mammalian contexts preferentially upregulate the first pathway enzyme PHGDH^66^, Yki tumors more prominently increase expression of the downstream enzymes (for example, *Psat* and *aay*) and expand both labeled and unlabeled serine pools, indicating conservation of pathway output with divergence in regulatory entry points across tumor contexts.

Additional nodes of the Yki metabolic network align with emerging themes from mammalian cancer metabolism. In human tumors, the G3P shuttle can couple glycolytic NAD⁺ regeneration to lipid synthesis and/or mitochondrial redox signaling^32,67^. Yki tumors similarly show increased glucose-derived G3P labeling with increased *Gpo1* expression, consistent with engagement of this shuttle to support high glycolytic flux and redox balance. In many cancers, the TCA cycle is reorganized around α-KG, which functions as a biosynthetic and regulatory branch point rather than a purely oxidative intermediate^68^. In Yki tumors, [U-¹³C₆]glucose tracing indicates that α-KG acts predominantly as a drain node, channeling glucose-derived carbon into glutamate, glutamine, aspartate and 2-HG under glucose-replete conditions, consistent with a conserved role in meeting biosynthetic and redox demands. Finally, whereas some cancers such as leukemia increase PPP activity to supply R5P and NADPH, Yki tumors show a bias toward increased non-oxPPP carbon rearrangements, moderate enrichment of labeled R5P and substantial glucose contribution to pyrimidine nucleotides. This pattern resembles settings such as colorectal cancer, where non-oxPPP enzymes including transketolase have been linked to growth and metastasis^69^, despite the absence of strong *Tkt* upregulation in this model. Collectively, these cross-species parallels support a conserved strategy in which glucose-derived carbon is redistributed into serine, glutamate/glutamine, aspartate and nucleotide synthesis to support proliferative growth.

*Drosophila melanogaster* therefore provides a complementary in vivo platform for defining conserved principles of tumor metabolism. Core metabolic pathways and oncogenic signaling modules relevant to human cancers are retained, while reduced tissue complexity, short generation times and genetic tractability enable systematic perturbation in intact organisms. Tissue-specific, temporally controlled genetic tools allow precise modeling of oncogenic events and interrogation of metabolic gene function under defined nutritional conditions^12,13^. Within this framework, our study establishes a tractable model for Hippo–YAP/TAZ/Yki-driven tumor metabolism. Hippo–YAP/TAZ signaling integrates mechanical, nutritional and mitogenic cues with metabolic state^22,23,70^, and YAP/TAZ hyperactivation contributes to epithelial tumorigenesis and therapy resistance, however, how Hippo pathway dysregulation reshapes central carbon handling at organismal scale has remained incompletely defined. Our current data begin to address this gap by mapping coordinated changes in glucose utilization in vivo following Yki activation.

Although multiple *Drosophila* tumor models have been generated through oncogene activation or tumor-suppressor inactivation, comparatively few studies have examined in vivo central carbon rewiring at a subset of metabolic reactions, or compared these programs with their mammalian tumor counterparts. This work addresses this gap by providing a pathway-level view of how glucose-derived carbon is redistributed across glycolysis, the TCA cycle, metabolic shuttles and the PPP in Yki-driven gut tumors. The resulting reference framework highlights branch points at which carbon is preferentially diverted toward biosynthetic or redox-supporting fates, thereby prioritizing metabolic nodes for functional genetic interrogation in a whole-animal context. Alignment with mammalian phenotypes further supports the relevance of these principles beyond *Drosophila* physiology. Together, these comparisons demonstrate that central carbon rewiring is not an idiosyncratic, mutation-specific phenomenon but a systematic, conserved form of metabolic adaptability that oncogenic signaling pathways exploit.

Finally, the multi-omic integration used here underscores the importance of resolving central carbon metabolism across regulatory layers. Transcript changes alone did not uniformly predict flux and pathway routing, and metabolite abundances could obscure shifts in labeled carbon flow, emphasizing that interpretations based on single modality can be incomplete or misleading. In mammalian systems, in vivo isotope tracing and multi-omic profiling have begun to define tumor nutrient use, but interpretation is often complicated by tissue heterogeneity and microenvironmental variability; for example, human NSCLC isotope-infusion studies reveal substantial intra- and inter-tumor heterogeneity linked to perfusion, and mouse lung tumor models show that in vivo nutrient usage and pathway requirements can diverge from cultured cells^58,71^. In contrast, the Yki fly tumor model offers genetic and nutritional control that should enable extension of the current “carbon fate map” toward quantitative flux-resolved analyses using staged tracer designs (for example, ¹³C-glucose and ¹³C-glutamine infusions). This multi-omic framework therefore provides a foundation for flux-resolved testing which central-carbon branchpoints are rate-limiting, which are governed primarily by enzyme abundance versus allostery, and which rewiring events are most conserved across oncogenic contexts and species.

## Materials and Methods

### Fly stocks and maintenance

Yki tumors were induced in *Drosophila* adult midgut intestinal stem cells using UAS-Yki^3SA^ (*w*;; UAS-yki.S111A.S168A.S250A.V5*; BDSC #228817) crossed to EGT (*tub-GAL80^ts^, esg-GAL4, UAS-GFP (II*)), following previously described procedures^29^. Briefly, crosses were set up using virgin female EGT flies and male UAS-Yki3SA flies and maintained for 3–4 days at 18°C. Flies were then transferred to new bottles to expand the cross. At 18°C, GAL80^ts^ is active and suppresses Gal4-mediated transgene expression. Adult progenies were collected within 3 days after eclosion, mated at 18°C for an additional 2 days, and then male adults were shifted to 29°C to induce transgene expression (e.g., “Day 8” indicates 8 days of Yki^3SA^ induction at 29°C). Progeny from a cross between EGT and w^11^^18^ were used as controls. During incubation at 29°C, flies were transferred to fresh food every 2 days until sample collection. Male flies were used in all experiments to minimize variability associated with female reproductive physiology and steroid hormone/ecdysone fluctuations. All flies were maintained on standard cornmeal–yeast–agar medium at 18°C or 25°C under a 12-hour light/dark cycle, unless otherwise noted. Standard laboratory food was prepared with the following composition: 12.7 g/L deactivated yeast, 7.3 g/L soy flour, 53.5 g/L cornmeal, 0.4% agar, 4.2 g/L malt, 5.6% corn syrup, 0.3% propionic acid, and 1% Tegosept/ethanol.

### Immunofluorescence and confocal imaging

Adult midguts were dissected in ice-cold 1× PBS and fixed in 4% paraformaldehyde in PBS for 30 min at room temperature. Samples were then rinsed and incubated for 1 h in 1× PBST (PBS containing 0.1% BSA and 0.3% Triton X-100) supplemented with 5% normal donkey serum (NDS) for permeabilization and blocking. Nuclei were counterstained with DAPI.

For phospho-histone H3 (pH3) staining, midguts were blocked in 5% NDS in PBST and incubated overnight at 4°C with rabbit anti-pH3 (1:500, CST 9701L) diluted in PBST. The following day, tissues were washed and incubated with donkey anti-rabbit Alexa Fluor 594 (1:1,000, Invitrogen A12381) for 1 h at room temperature.

After three washes in PBST, guts were mounted in Vectashield with DAPI (Vector Laboratories, H-1200).

For BODIPY lipid staining, guts were dissected and fixed as described above, washed three times in 1× PBS, and incubated with BODIPY 493/503 (Invitrogen D3922; 1 mg/mL in 1× PBS) for 30 min at room temperature. Samples were then washed three times in PBS and mounted in Vectashield with DAPI.

Confocal images were acquired using a Zeiss LSM980 microscope equipped with Airyscan 2 and ZEN software at the Microscopy Resources on the North Quad (MicRoN) core facility at Harvard Medical School. pH3-positive cells were quantified by manually counting labeled nuclei using Fiji/ImageJ.

### Lifespan assay

Lifespan was assessed by monitoring the survival of flies under conditions with or without Yki tumor induction. For assays with Yki tumors, flies were incubated at 29°C; for assays without Yki tumors, flies were maintained at 18°C. Male flies were placed in vials containing fresh food, and the number of dead flies was recorded daily. For each lifespan experiment at 29°C, 20 flies were used per vial (replicate), with a total of 5 biological replicates. Flies were transferred to fresh food every 2 days until the end of the experiment.

### Polar metabolites extraction and LC-MS profiling

Intracellular metabolites were extracted using 80% (v/v) aqueous methanol prepared in HPLC-grade water. For each replicate, 65 adult *Drosophila* midgut tissues (Yki tumors and controls) were dissected in ice-cold 1× PBS and transferred to a 1.5 mL Eppendorf tube containing 400 µL ice-cold 80% HPLC-grade methanol and 0.5 mm zirconium beads. Tissues were homogenized using a Bullet Blender tissue homogenizer (model BBX24, Next Advance) for three cycles of 5 min at speed 9, assessing homogenate uniformity between cycles.

Following homogenization, 5 µL of each sample was transferred to a new 1.5 mL Eppendorf tube for subsequent BCA protein quantification. The remaining homogenate was centrifuged at 20,000 × g for 5 min at 4°C, and the supernatant was transferred to a new 1.5 mL Eppendorf tube. To maximize extraction efficiency, the pellet was resuspended in an additional 300 µL of 80% methanol, subjected to the same homogenization and centrifugation conditions, and the resulting supernatant was combined with the initial supernatant.

Combined extracts were dried by SpeedVac centrifugation and stored at −80°C until mass spectrometric analysis.

Prior to analysis, metabolite pellets were resuspended in 22 µL HPLC-grade water. Metabolomics data were acquired by liquid chromatography–tandem mass spectrometry (LC-MS/MS) as previously described^72^ at the Beth Israel Deaconess Medical Center Mass Spectrometry Facility. Briefly, 8 µL of each resuspended sample was injected and analyzed on a hybrid 6500 QTRAP hybrid triple quadrupole mass spectrometer (SCIEX) coupled to a Prominence UFLC HPLC system (Shimadzu) for steady-state metabolite profiling. Selected reaction monitoring (SRM) was used to quantify approximately 310 polar metabolites, employing positive/negative ion polarity switching with hydrophilic interaction liquid chromatography (HILIC) (Amide Xbridge column, Waters). Q3 peak areas from each metabolite SRM transition were integrated using MultiQuant v3.2 software (SCIEX) and obtained results were normalized to total protein abundance as determined by BCA assay.

Total protein-normalized intensities were processed: first, filtered for missingness. Metabolites were retained if at least one of the two conditions exhibited <20% missing values. Remaining metabolites were mapped to KEGG IDs, and those without KEGG annotations were removed. For metabolites with duplicate entries, the entry with the higher mean intensity across all samples was retained. Intensities were log-transformed prior to imputation. Missing values were imputed using QRILC (Quantile Regression Imputation of Left-Censored data) implemented in the impute.QRILC function from the imputeLCMD package (v2.1), which draws random values from an estimated left-censored distribution based on quantile regression. The resulting imputed data matrix was uploaded to the MetaboAnalyst web platform, and the Enrichment Analysis module was used to obtain pathway enrichment scores.

For relative abundance analysis and figure generation of individual metabolites, the imputed data matrix was normalized to the median value of each sample. This transformation yields a value of 0 for metabolites with intensities near the sample median, negative values for metabolites with intensities below the median, and positive values for those above the median. Statistical analysis was performed using MetaboAnalystR^73^ and ggplot2 package was used for comparison of the means of the normalized metabolite peaks in R.

### [U-¹³C₆]glucose isotope tracing

For targeted isotope tracing, flies were first starved on distilled water for 8 h at 29°C to empty the gut prior to ¹³C-labeled glucose feeding. Flies were then transferred to [U-¹³C₆]glucose tracer mixed with blue dye and fed for ∼8 h at 29°C. The presence of blue dye in the gut confirmed tracer consumption.

Gut tissues were dissected, and intracellular metabolites were extracted and prepared for mass spectrometric analysis as described above. Q1/Q3 SRM transitions for ¹³C incorporation into polar metabolite isotopomers were established, and data were acquired by LC-MS/MS via SRM^74^ at the Beth Israel Deaconess Medical Center Mass Spectrometry Facility. Peak areas were generated for each detected isotopomer using MultiQuant (version 2.1) software and normalized to total protein abundance as measured by BCA assay.

### Long-read bulk RNA sequencing and data analysis

Three biological replicates of day 8 *Drosophila* midguts of Yki tumors and control were dissected in ice-cold 1X PBS, and total RNA was extracted using the Direct-zol MniPrep kit (Zymo Research, Cat. R2050) according to the manufacture’s instructions. RNA quality was assessed using Agilent TapeStation High Sensitivity RNA assay at the Biopolymer Facility (BPF), Harvard Medical School. Samples with RNA integrity number (RIN) > 9.6 were useded for cDNA library preparation.

cDNA libraries were generated using the Oxford Nanopore Technology (ONT) cDNA-PCR Barcoding Kit v14 (SQK-PCB114.24), following the manufacturer’s protocol with minor adjustment to accommodate the input RNA volume. A total of 1.5ug of total RNA per sample was used, and reaction volumes were scaled up accordingly. The resulting libraries contained full-length cDNA suitable for long-read nanopore sequencing. Library quality was evaluated using Qubit fluorometry and Agilent TapeStation High Sensitivity D5000 assay at the BPF. Prepared cDNA libraries were sequenced on an ONT PromethION flow cell at the Bauer Sequencing Core, Harvard University.

Raw nanopore reads were trimmed and filtered for low-quality sequences using Pychopper v2, and the processed reads were aligned to the *Drosophila melanogaster* reference genome (BDGP6.46) using minimap2. Transcript and gene-level counts were quantified with Bambu (v3.10.1). Differential gene expression analysis was performed in R (v4.3.2) using the standard DESeq2 (v1.48.2) workflow for normalization and testing.

Gene group enrichment of differentially expressed genes (DEGs) was performed using Gene List Annotation for Drosophila (GLAD)^75^ and results were visualized in R (v4.3.2) and ggplot2 package was used for comparison of the means of the gene expression.

### Human cancer dataset retrieval and processing

The human cancer datasets were retrieved from the data made available for Benedetti et al. (2023)^76^ from the Reznik lab at https://zenodo.org/records/7348648. Version 0.3.2 of the pancancer metabolomics data was downloaded to use as the human cancer raw data, containing both metabolomics and transcriptomics datasets. The metabolomics datasets had already been preprocessed by Benedetti et al., as described in their paper. Additionally, for each cancer type, metabolites were mapped to KEGG IDs and unmapped metabolites were filtered out. The variance of duplicate mapping KEGG IDs were calculated and compounds with the lowest variance in normal samples were maintained. Each of the processed datasets were then input to Metaboanalyst, using the Statistical Analysis module to calculate differential expression of metabolites and the Enrichment Analysis module for pathway enrichment. The intensity data was then normalized by log10 transformation, and the metadata, normalized intensity data, and metabolite differential expression output by Metaboanalyst were combined for each cancer type to make the master data frame.

### Statistical methods

The statistical tests used for the individual analyses are indicated in the figure legends or corresponding method descriptions. Generally, two-tailed t tests were used for comparison between two groups using GraphPad Prism (v11.0.0) or R (R 4.5.2). Reported p-values for individual metabolite or gene comparisons were not adjusted for multiple hypothesis testing.

## Data availability

All data generated in this study including metabolomics, 13C-glucose isotope tracing and long-read sequencing are provided in the Source Data. Processed and analyzed data are provided in Supplementary Data.

## Acknowledgements

The authors thank R. Binari, S. Mohr, B. Mathey-Prevot and all members of the Perrimon laboratory for their critical insights and assistance throughout this research. We are grateful to the Reznik laboratory for generously sharing human cancer datasets and providing guidance on data analysis. We thank K. Rahimi and W. Chen for assistance with the initial nanopore sequencing setup, and The Bauer Core Facility at Harvard University for support with nanopore sequencing. We also thank P.M. Llopis and the Microscopy Resources on the North Quad (MicRoN) core facility at Harvard Medical School for support with confocal imaging, and the Biopolymer Facility at Harvard Medical School for RNA/DNA quality assessment. This work is funded by NIH 5R01DK136945 and the CANCAN team supported by the Cancer Grand Challenges partnership funded by Cancer Research UK (CGCATF-2021/100022) and the National Cancer Institute (1 OT2 CA278685-01). N. Perrimon is an investigator of the Howard Hughes Medical Institute. This article is subject to HHMI’s Open Access to Publications policy.

## Author information

### Authors and Affiliations

**Department of Genetics, Blavatnik Institute, Harvard Medical School, Boston, MA, USA**

Younshim Park

Mujeeb Qadiri

Yanhui Hu

Norbert Perrimon

**Division of Signal Transduction, Beth Israel Deaconess Medical Center, Boston, MA, USA**

John M. Asara

**Howard Hughes Medical Institute, Boston, MA, USA**

Norbert Perrimon

## Contributions

Y.P. and N.P. conceived the study and designed the experiments. Y.P. conducted the majority of the experiments and the data analysis. J.A. conducted metabolomics and [U-¹³C₆]glucose tracing measurements. M.Q., Y.H. conducted the bioinformatics analysis. Y.P. and N.P. interpreted the results and wrote the manuscript with contributions and feedback from all authors. All authors reviewed and approved the final manuscript.

## Corresponding authors

Correspondence to Younshim Park or Norbert Perrimon

---

**Figure S1.**
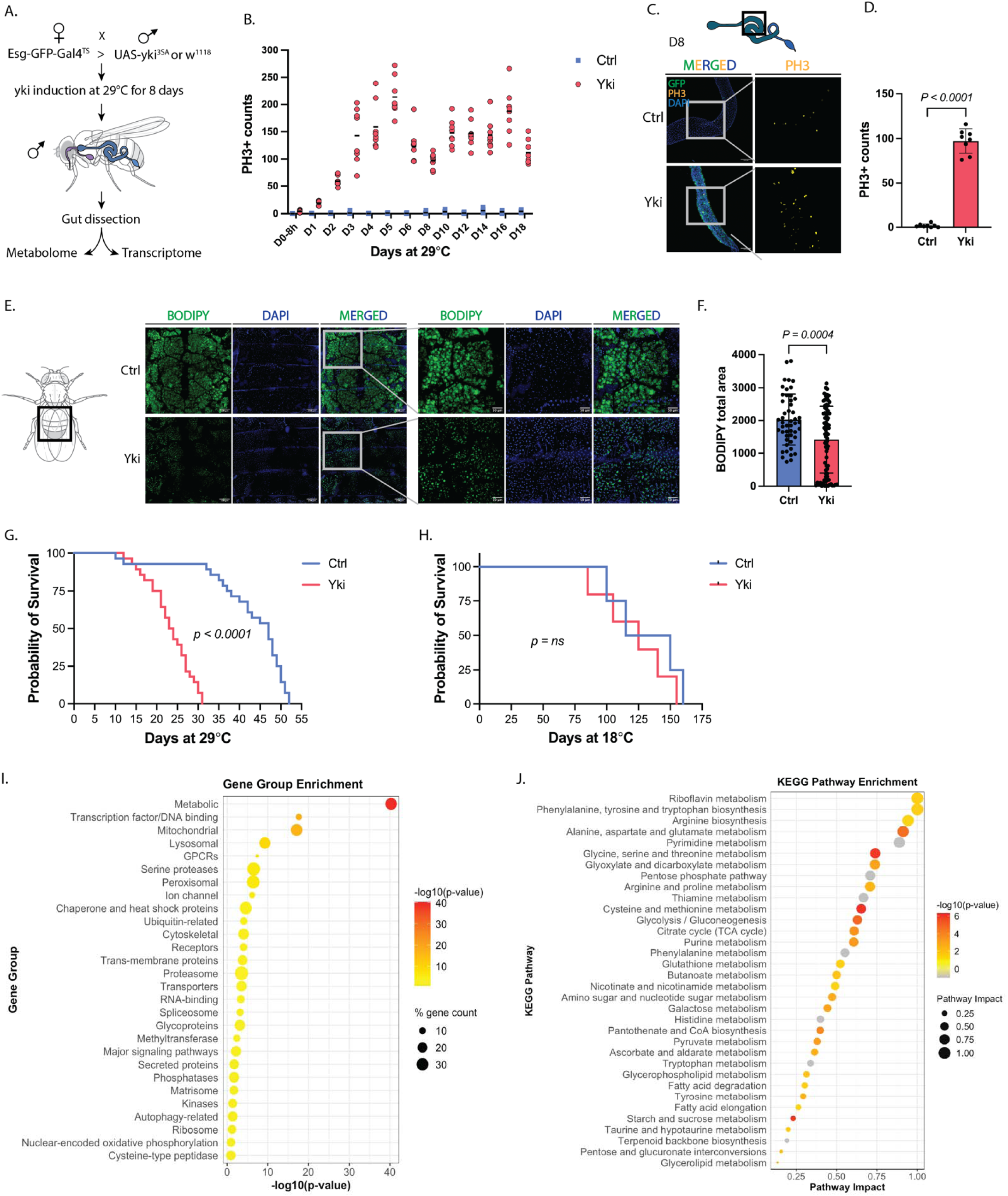
Yki tumors recapitulate organismal hallmarks associated with human cancers. (A) Experimental schematic for Yki tumor induction and sample collection. yorkie (yki^3SA^) was overexpressed in intestinal stem cells using a temperature-sensitive gene-expression driver; flies were shifted to 29°C for 8 days, followed by gut dissection for metabolome and transcriptome profiling. (B) Time-course quantification of PH3-positive (PH3+) cells during induction at 29°C demonstrating sustained proliferative activity over time. PH3+ cells were counted in whole Yki guts. (C-D) Representative images of phospho-histone H3 (PH3) immunostaining in control and Yki-induced midguts at day 8 (C) and quantification of PH3+ cells (D), indicating elevated mitotic activity and increased proliferation in Yki tumors. (E-F) Representative BODIPY staining of control and Yki tumor-bearing flies (E) and quantification of BODIPY signal (F) showing decreased lipid abundance in Yki flies, consistent with a cachexia-like phenotype. (G-H) Lifespan analysis at 29°C (G) and under non-inducing control conditions (18°C) (H) showing shortened survival upon yki^3SA^induction at 29°C, whereas no significant survival difference is observed between genotypes at 18°C. (I) Gene group enrichment of differentially expressed genes in Yki tumors highlighting significant enrichment of the metabolic gene group. (J) Pathway enrichment analysis of metabolite profiling data showing broad metabolic remodeling, with prominent alterations in bioenergetic and macromolecule-related pathways, including central carbon metabolism, amino acid metabolism and nucleotide metabolism.

**Figure S2.**
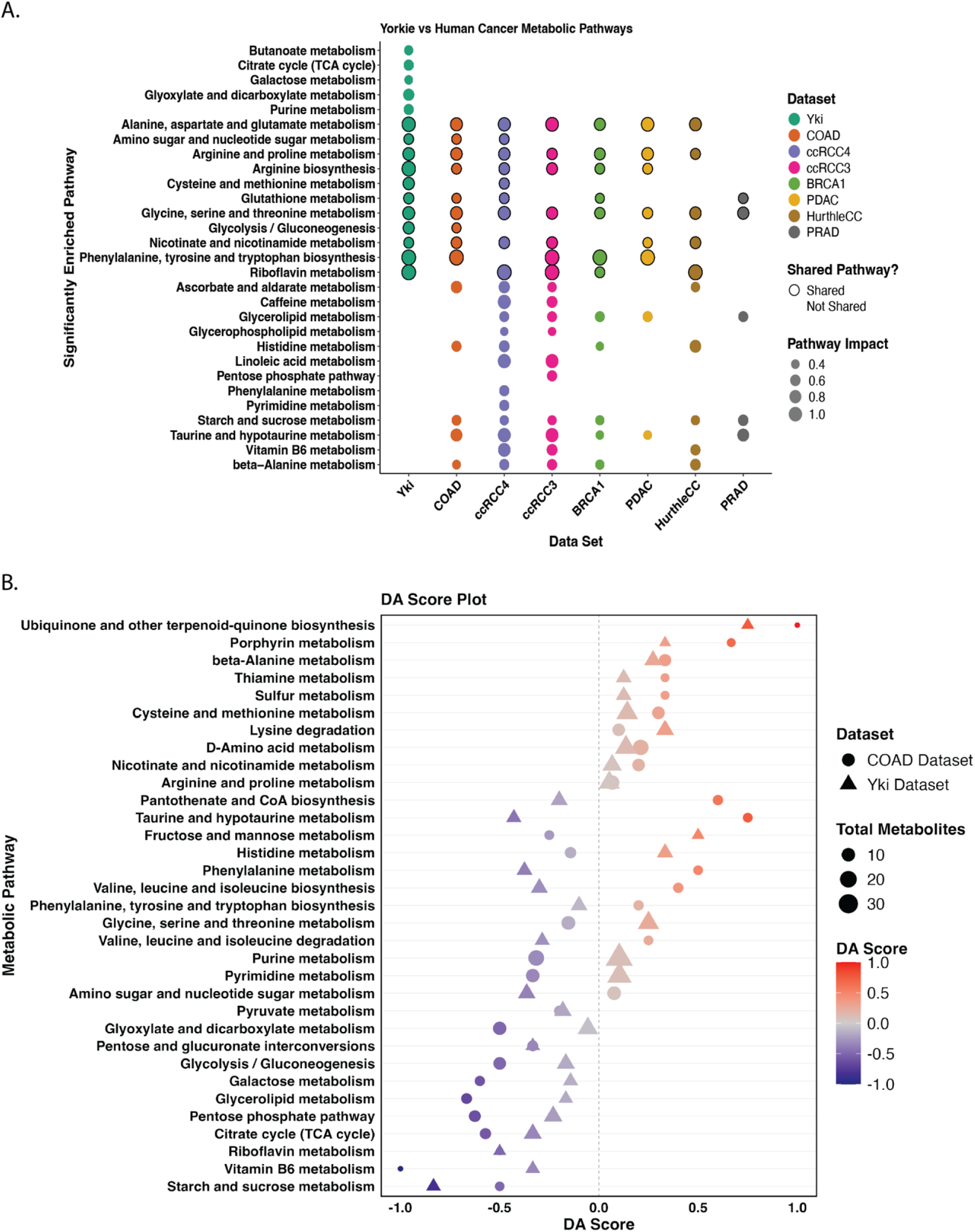
Cross-species comparison reveals shared metabolic pathway rewiring in Yki and human tumors. (A) Pathway enrichment analysis of Day 8 Yki tumor metabolomics compared with six human cancer types plus one additional subtype. Sixteen KEGG pathways with pathway impact > 0.4 were selected and compared across seven high-quality human cancer metabolomics datasets. Eleven of the 16 Yki-enriched pathways are also enriched in multiple human cancers, with the strongest metabolic overlap observed with human colon cancer (COAD) and clear cell renal cell carcinoma subtype 4 (ccRCC4). (B) Differential Abundance (DA) analysis comparing Yki tumors and human colon cancer (COAD) reveals concordant up-or downregulation in approximately two-thirds of shared pathways, including glycolysis and amino acid metabolism. The DA score summarizes pathway directionality and is calculated as (number of upregulated metabolites – number of downregulated metabolites)/total number of significant metabolites in the pathway, ranging from +1 (all upregulated) to -1 (all downregulated), with 0 indicating either no significant metabolites or balanced up- and downregulation.

**Figure S3 (Related to Figure 1).**
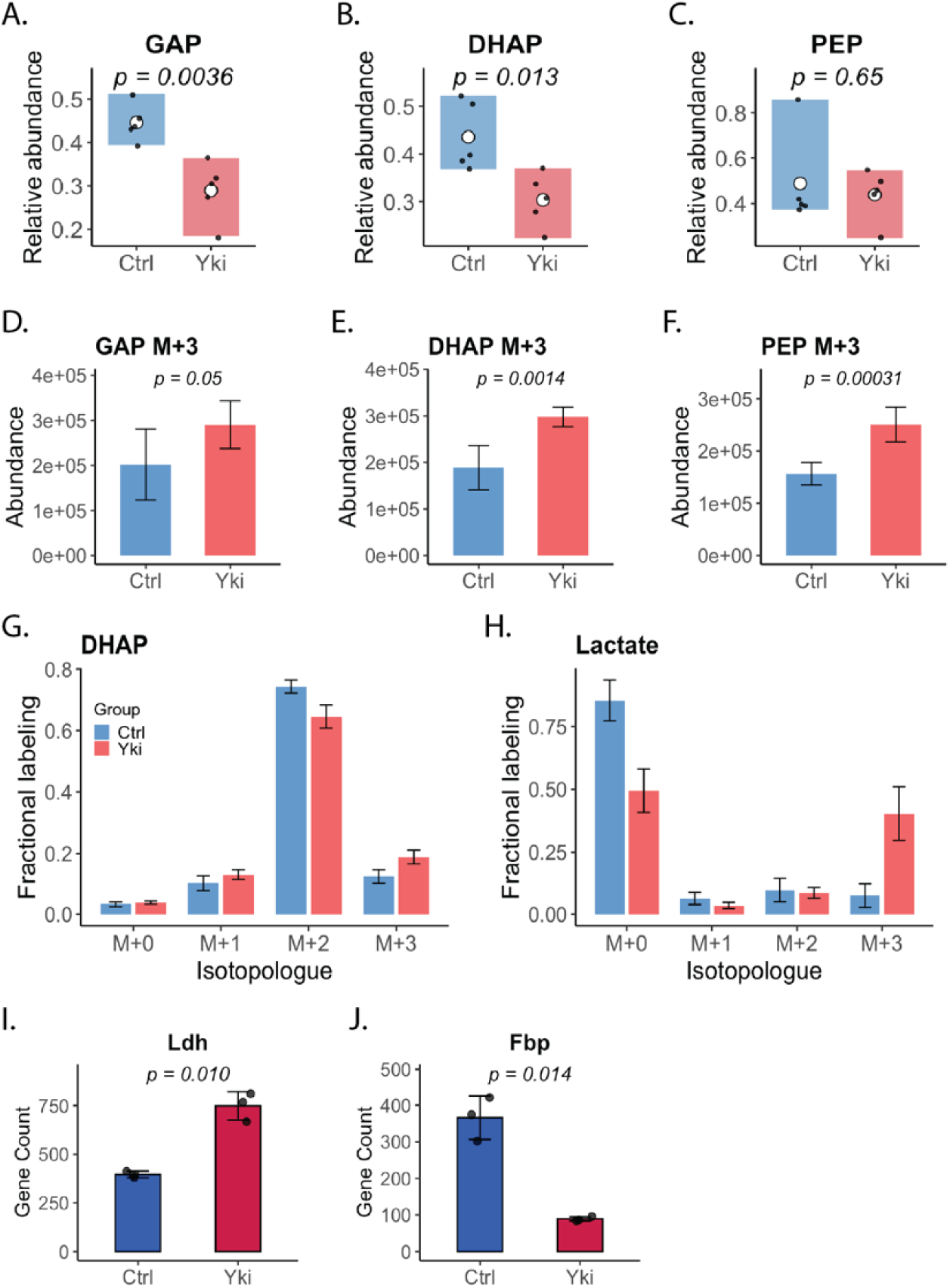
Expanded metabolic evidence for enhanced glycolytic flux and preferential routing of glucose-derived carbon to lactate in Yki tumors. (A-C) Relative abundance of the lower-glycolytic intermediates GAP, DHAP and PEP. (D-F) [U-¹³C₆]glucose tracing showing increased abundance of the fully labeled M+3 isotopologues of GAP, DHAP and PEP in Yki tumors, consistent with enhanced propagation of glucose-derived carbon through lower glycolysis. (G-H) Fractional labeling of DHAP isotopolgue (M+0-M+3) (G) and lactate (M+0-M+3) (H). Lactate isotopologue fractional labeling shows a decreased in the unlabeled (M+0) fraction and an increased in M+3 lactate in Yki tumors, indicating enhanced incorporation of glucose-derived carbon into lactate. (I-J) Transcript abundance for Ldh (I) and Fbp (J), indicating increased expression of lactate dehydrogenase and reduced expression of fructose-bisphosphatase in Yki tumors. Reduced expression of Fbp and Pepck, which encode rate-limiting enzymes of gluconeogenesis, suggests that glucose-derived carbon flux is preferentially directed downstream through glycolysis rather than diverted into reverse gluconeogenic reactions.

**Figure S4 (Related to Figure 2).**
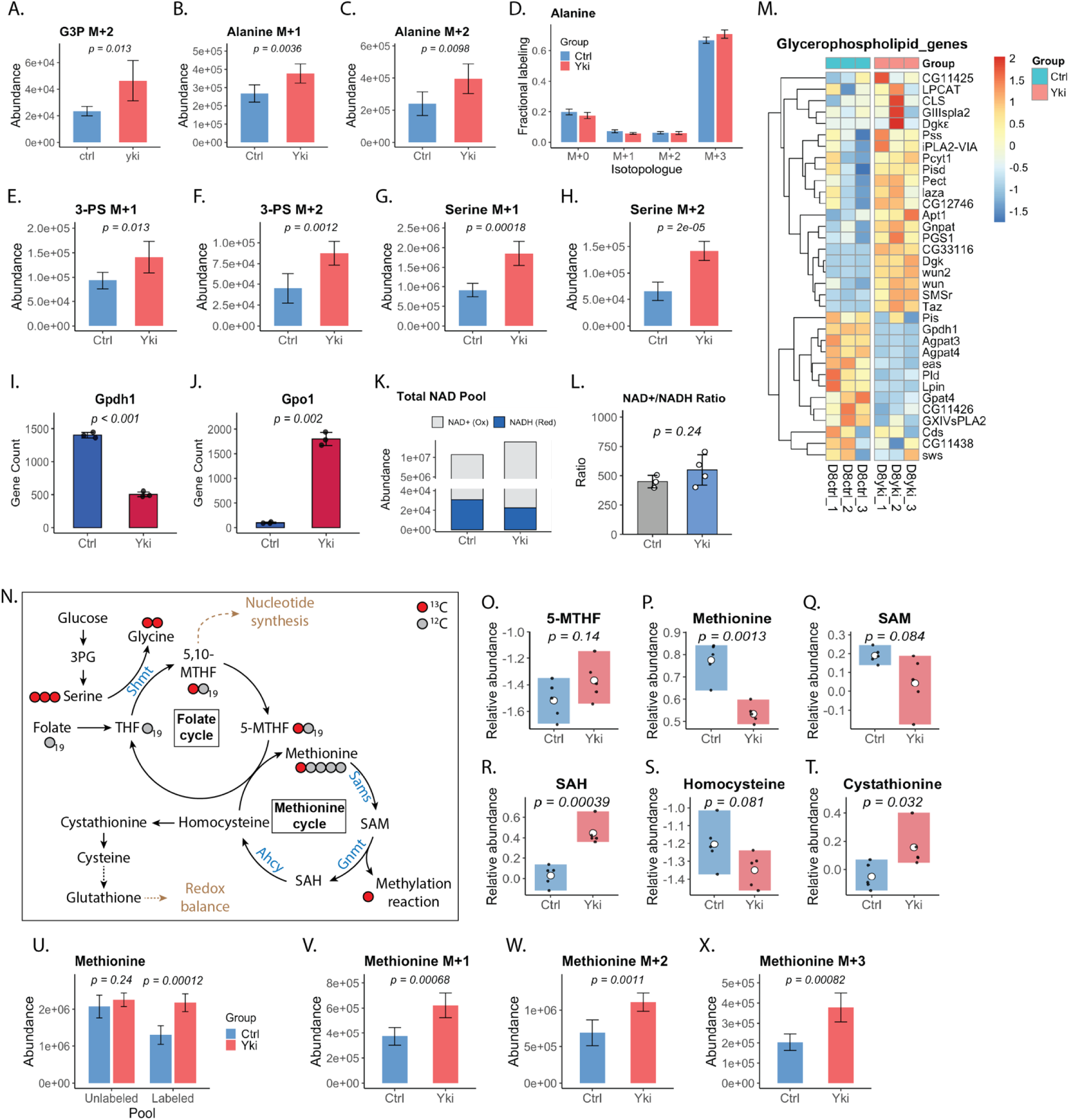
Expanded analysis of G3P shuttle activity, alanine/serine labeling, redox state, and one-carbon–methionine metabolism in Yki tumors. (A-C) Abundance of individual istopologues from [U-¹³C₆]glucose tracing in Ctrl and Yki samples. Abundances of G3P M+2 (A), alanine M+1 (B), and M+2 (C). (D) Fractional labeling of alanine isotopologues (M+0–M+3). (E-H) Abundances of 3-PS M+1 (E) and M+2 (F), and abundances of serine M+1 (G) and M+2 (H) (I, J) RNA-seq expression of Gpdh (I; cytosolic glycerol 3-phosphate dehydrogenase) and Gpo1 (J; mitochondrial G3P dehydrogenase) in Ctrl and Yki samples, showing normalized gene counts per sample. (K) Total NAD pool measured by LC–MS, plotted as stacked abundances of oxidized (NAD⁺) and reduced (NADH) species. (L) NAD⁺/NADH ratio in control and Yki samples.(M) Heatmap of transcripts encoding enzymes involved in glycerophospholipid metabolism. (N) Schematic of the folate and methionine cycles linking glucose-derived serine to glycine, one-carbon units, methionine, S-adenosylmethionine (SAM), S-adenosylhomocysteine (SAH), and transsulfuration to cystathionine, cysteine, and glutathione; red circles denote potential incorporation of ^13C from [U-¹³C₆]glucose. (O-T) Relative abundances of 5-methyltetrahydrofolate (5-MTHF; O), methionine (P), SAM (Q), SAH (R), homocysteine (S), and cystathionine (T) in control and Yki samples. (U) Abundances of unlabeled and total ¹³C-labeled methionine pools following [U-¹³C₆]glucose tracing. (V–X) Abundances of methionine M+1 (V), M+2 (W), and M+3 (X) isotopologues, reflecting incorporation of one, two, or three ¹³C atoms, respectively.

**Figure S5 (Related to Figure 3).**
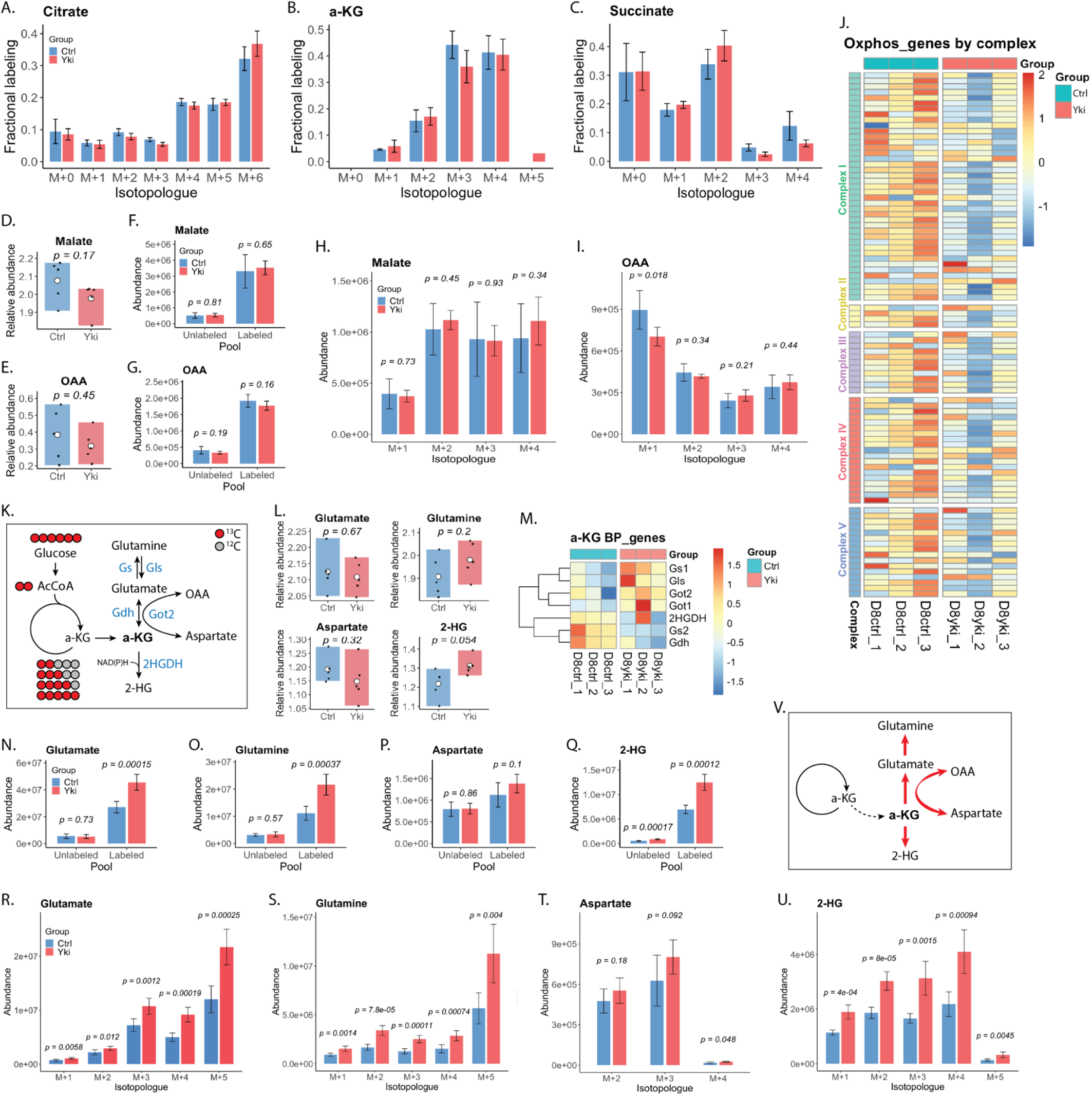
Glucose-derived carbon is preferentially partitioned at the α-KG node into biosynthetic branches in Yki tumors. (A–C) Fractional ¹³C labeling distributions for citrate, α-KG and succinate, highlighting strong labeling in upstream metabolites and an isotopic discontinuity between α-KG and succinate.(D–G) Additional measurements of downstream TCA-associated intermediates and pools (malate and OAA), including unlabeled versus labeled pool comparisons where indicated. (H–I) Malate and OAA isotopologue abundances (M+1–M+4), providing further support for preserved glucose contribution to late intermediates despite altered TCA organization. (J) Heatmap of oxidative phosphorylation (OxPhos) gene expression grouped by respiratory complex, showing coordinated remodeling of mitochondrial bioenergetic programs in Yki tumors. Genes assigned to each complex are listed in the Supplementary Table 1. (K) Schematic of α-KG branch pathways connecting the TCA cycle to synthesis of glutamate/glutamine, aspartate, and 2-HG. (L) Relative abundances of α-KG-connected branch metabolites (glutamate, glutamine, aspartate and 2-HG). (M) Heatmap of transcripts encoding α-KG branch-point enzymes (e.g., aminotransferases, glutamine synthetase and related nodes), supporting increased capacity for α-KG exit flux. (N–Q) Unlabeled versus ¹³C-labeled pool abundances for glutamate, glutamine, aspartate and 2-HG, showing increased glucose-derived labeling of α-KG-connected products in Yki tumors. (R–U) Isotopologue abundances for glutamate, glutamine, aspartate and 2-HG. Notably, M+5 glutamate/glutamine is readily detected and increased in Yki tumors despite undetectable α-KG M+5 in the bulk pool (see panel B in the main figure set), consistent with a rapidly turning-over, highly labeled α-KG sub-pool feeding amino-acid synthesis. (V) Summary schematic illustrating preferential routing of α-KG-derived carbon into biosynthetic outputs (glutamate/glutamine/aspartate and 2-HG) in Yki tumors.

**Figure S6 (Related to Figure 4).**
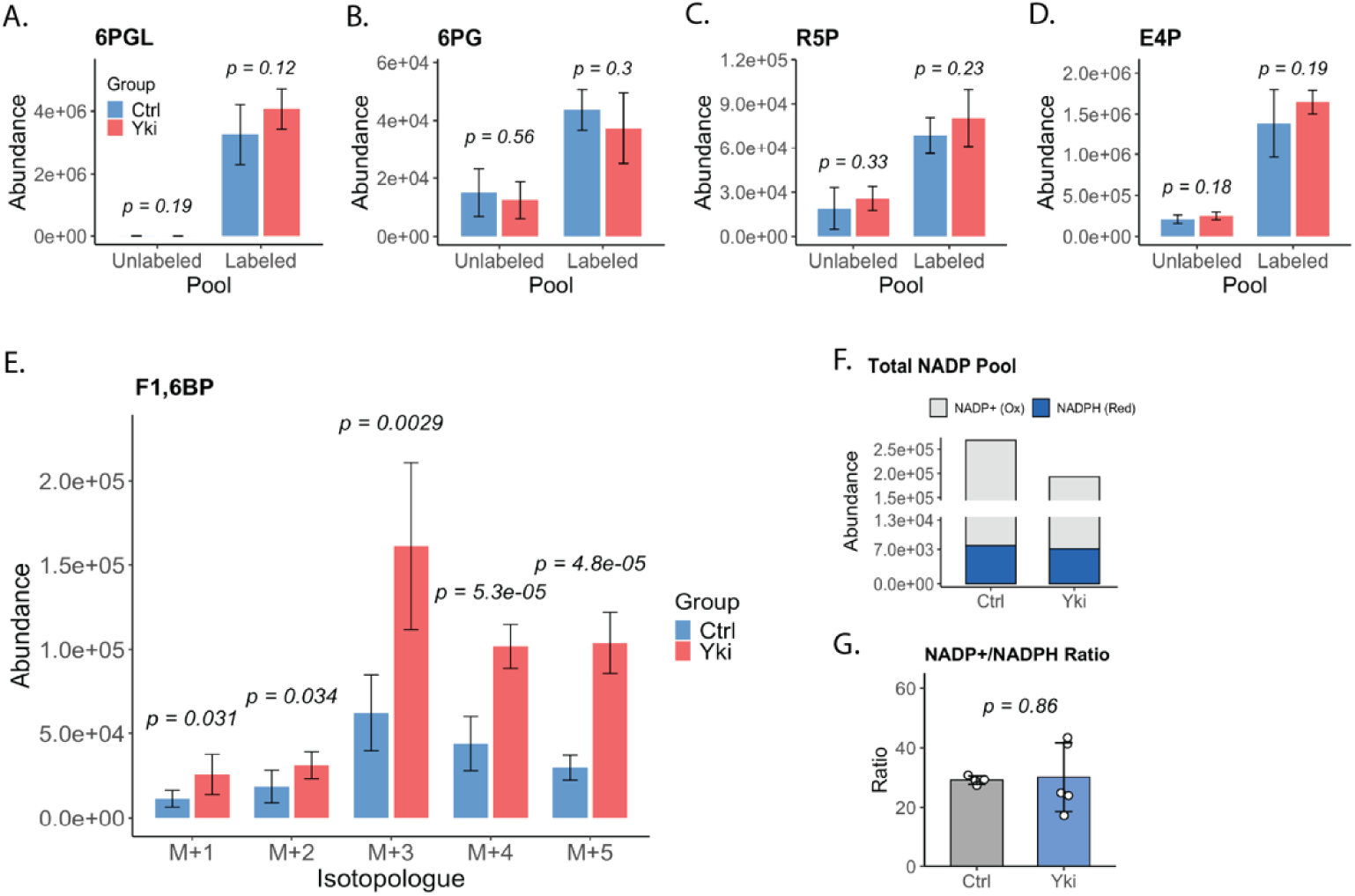
Expanded metabolic evidence for enhanced non-oxidative PPP activity and preserved NADP(H) redox state in Yki tumors. (A–C) Abundance of unlabeled and [U-¹³C₆]glucose-derived (labeled) intermediates of the oxidative PPP (oxPPP): 6-phosphogluconolactone (6PGL), 6-phosphogluconate (6PG), and ribose-5-phosphate (R5P). Labeled species dominate each pool, indicating strong glucose contribution, but total labeled + unlabeled pools are not significantly altered in Yki tumors relative to controls. (D) Abundance of unlabeled and labeled erythrose-4-phosphate (E4P), a non-oxidative PPP (non-oxPPP) intermediate, showing a trend toward increased labeled E4P in Yki tumors. (E) Abundances of individual fructose-1,6-bisphosphate (F1,6BP) isotopologues (M+1–M+5). Multiple partially labeled species are significantly enriched in Yki tumors, consistent with increased carbon shuffling through the non-oxPPP and reintegration of PPP-derived carbons into upper glycolysis. (F) Total NADP(H) pool resolved into oxidized NADP+ (grey) and reduced NADPH (blue). Yki tumors exhibit a selective decrease in NADP+ with relatively preserved NADPH levels, indicating a modest contraction of the overall NADP(H) pool while maintaining reducing capacity. (G) NADP+/NADPH ratio, showing no significant difference between control and Yki tumors, indicating that despite changes in pool size, the NADP(H) redox balance is maintained in Yki tumors.

**Figure S7 (Related to Figure 5).**
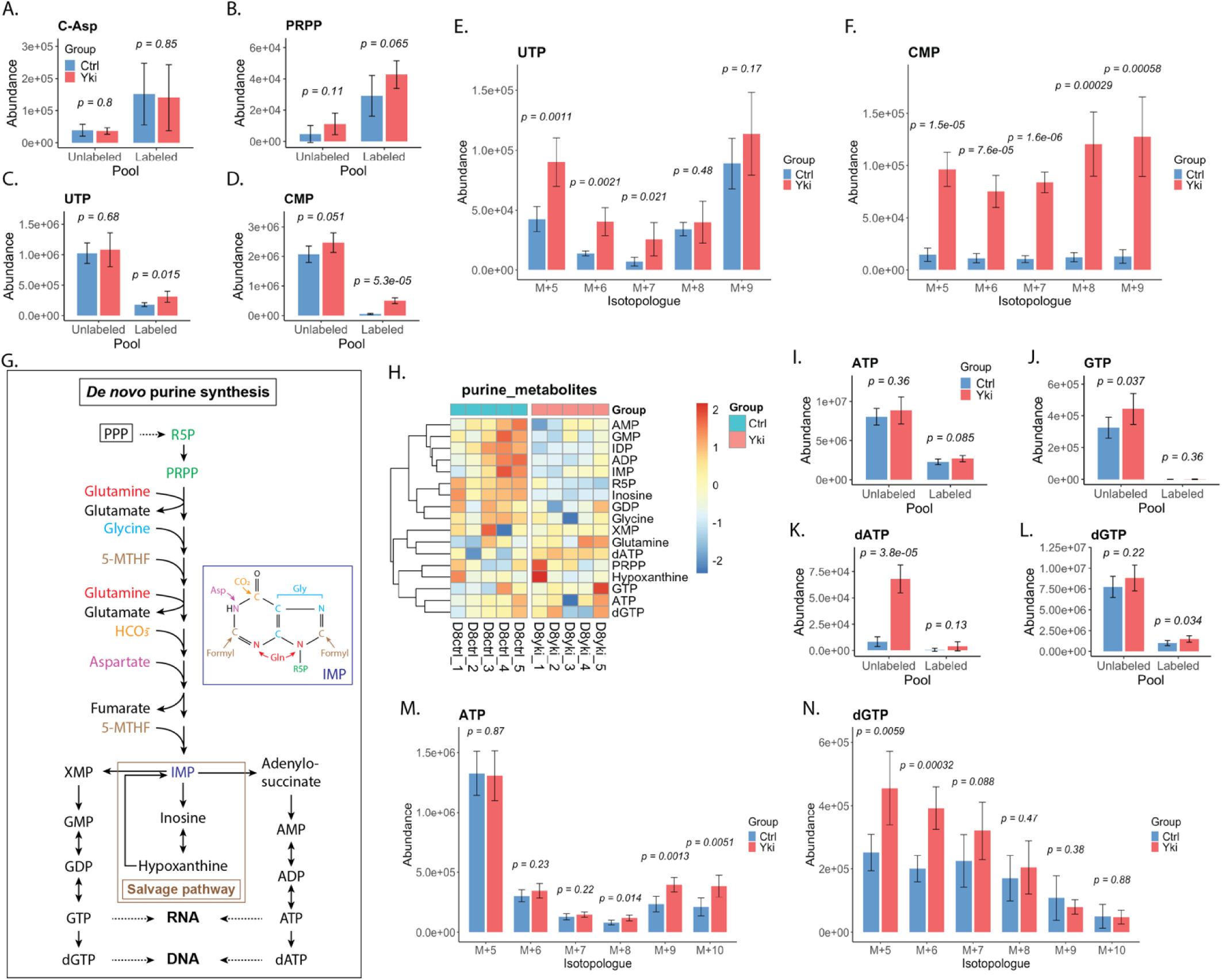
Glucose-derived carbon supports expanded pyrimidine and purine nucleotide pools in Yki tumors. (A–D) Abundance of unlabeled and ¹³C-labeled pools for selected nucleotide precursors and pyrimidine nucleotides (carbamoyl aspartate, PRPP, UTP, and CMP) following [U-¹³C₆]glucose tracing, indicating glucose contribution to nucleotide synthesis. (E–F) UTP and CMP isotopologue abundances (M+n), showing distribution of glucose-derived labeling across nucleotide carbon backbones. (G) Schematic of de novo purine synthesis and salvage. Enzymes are shown in blue and metabolites in black; major inputs (e.g., ribose/PRPP, glutamine, glycine, aspartate, bicarbonate/CO₂, and folate-derived one-carbon units) are indicated, with output into RNA- and DNA-directed nucleotide pools(H) Heatmap of purine metabolite abundances showing coordinated changes in adenine- and guanine-nucleotide pools in Yki tumors. (I–L) Abundance of unlabeled and ¹³C-labeled pools for ATP, GTP, dATP, and dGTP, reporting glucose contribution to ribose-containing purine nucleotides and deoxynucleotides. (M–N) ATP and dGTP isotopologue abundances (M+n), quantifying incorporation of glucose-derived carbon into purine nucleotide isotopologue pools.

**Supplementary Table 1.**
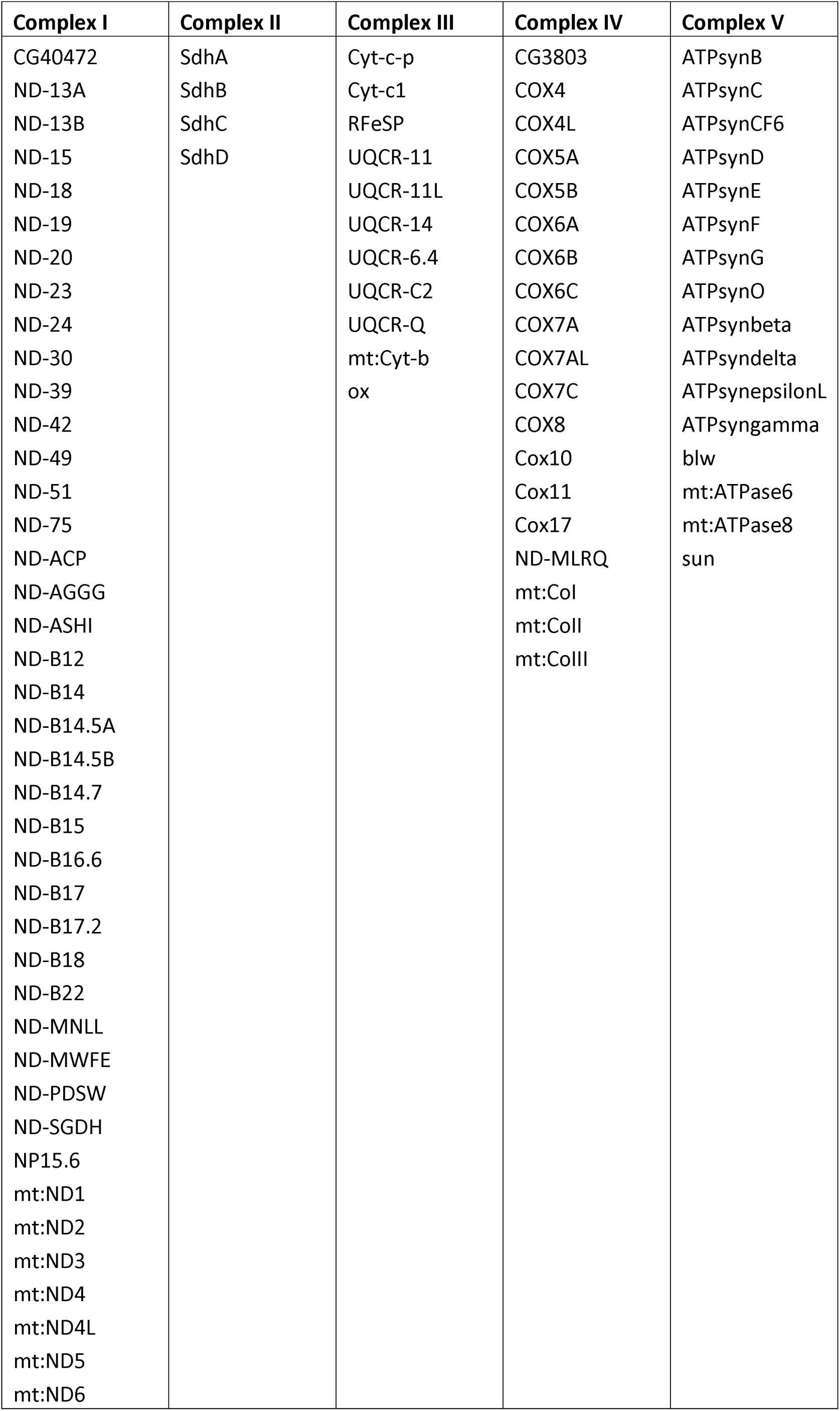
Oxidative phosphorylation (OxPhos) genes grouped by respiratory complex.

